# Active Listening

**DOI:** 10.1101/2020.03.18.997122

**Authors:** Karl J. Friston, Noor Sajid, David Ricardo Quiroga-Martinez, Thomas Parr, Cathy J. Price, Emma Holmes

**Author notes:** Address for correspondence: Emma Holmes, The Wellcome Centre for Human Neuroimaging, UCL Queen Square Institute of Neurology, London, UK WC1N 3AR.

## Abstract

This paper introduces active listening, as a unified framework for synthesising and recognising speech. The notion of *active listening* inherits from active inference, which considers perception and action under one universal imperative: to maximise the evidence for our (generative) models of the world. First, we describe a generative model of spoken words that simulates (i) how discrete lexical, prosodic, and speaker attributes give rise to continuous acoustic signals; and conversely (ii) how continuous acoustic signals are recognised as words. The ‘active’ aspect involves (covertly) segmenting spoken sentences and borrows ideas from active vision. It casts speech segmentation as the selection of internal actions, corresponding to the placement of word boundaries. Practically, word boundaries are selected that maximise the evidence for an internal model of how individual words are generated. We establish face validity by simulating speech recognition and showing how the inferred content of a sentence depends on prior beliefs and background noise. Finally, we consider predictive validity by associating neuronal or physiological responses, such as the mismatch negativity and P300, with belief updating under active listening, which is greatest in the absence of accurate prior beliefs about what will be heard next.

## Introduction

This paper could be read at three complementary levels: it could be regarded as a foundational paper introducing a *generative model* of spoken word sequences and an accompanying inversion (i.e., word recognition) scheme that has some biological plausibility; e.g., (Kleinschmidt and Jaeger 2015). Alternatively, one could read this article as a proposal for a speech recognition scheme based upon first (Bayesian) principles; e.g., (Rosenfeld 2000). Finally, one could regard this work as computational neuroscience, which makes some predictions about the functional brain architectures that mediate hierarchical auditory perception, when listening or repeating spoken words; e.g., (Hickok and Poeppel 2007, Houde and Nagarajan 2011, Tourville and Guenther 2011, Ueno, Saito et al. 2011). In the latter setting, the generative model can be used to predict the effects of synthetic lesions, i.e., as the basis for computational neuropsychology. In other words, one could optimise the parameters of the active listening scheme described below to best explain empirical (electrophysiological or behavioural) responses of individual subjects. We hope to pursue this in subsequent work. The current paper focuses on the form of the generative model, the accompanying recognition or inference scheme, and the kinds of behavioural and neuronal responses it predicts.

Speech recognition is not a simple problem. The auditory system receives a continuous acoustic signal and, in order to understand the words that are spoken, must parse a continuous signal into discrete words. To a naïve listener, the acoustic signal provides few cues to indicate where words begin and end (Altenberg 2005, Thiessen and Erickson 2013). Furthermore, even when word boundaries are made clear, there exists a many-to-many mapping between lexical content and the acoustic signal. This is because speech is not ‘invariant’ (Liberman, Cooper et al. 1967)—words are never heard out of a particular context. When considering how words are generated, there is wide variability in the pronunciation of the same word among different speakers (Hillenbrand, Getty et al. 1995, Remez 2010)—and even when spoken by the same speaker, pronunciation depends on prosody (Bänziger and Scherer, 2005). From the perspective of recognition, two signals that are acoustically identical can be perceived as different words or phonemes by human listeners, depending on their context—for example, the preceding words or phonemes (Mann 1980, Miller, Green et al. 1984), preceding spectral content (Holt, Lotto et al. 2000), or the duration of a vowel that follows a consonant (Miller and Liberman 1979). The current approach considers the processes involved in segmenting speech—and inferring the words that were spoken—as complementary.

The idea that speech segmentation and lexical inference operate together did not figure in early accounts of speech recognition. For example, the Fuzzy Logic Model of Perception (FLMP) (Oden and Massaro 1978, Massaro 1987, Massaro 1989) matches acoustic features with prototype representations to recognise phonemes, even when considered in the context of words and sentences. Similarly, the Neighbourhood Activation Model (NAM) (Luce 1986, Luce and Pisoni 1998) considers individual word recognition; it accounts for effects of word frequency, but does not address the segmentation problem. Later connectionist accounts, such as TRACE (McClelland and Elman 1986), assumed that competition between lexical nodes drives recognition, where competition is mediated by inhibitory connections between nodes: bottom-up cues determine recognition of phonemes and top-down cues take into account the plausible words in the lexicon. Shortlist B (Norris and McQueen 2008) reformulates this problem as one of an optimal Bayesian observer and incorporates word frequency effects.

Implicit in these connectionist and Bayesian accounts is the idea that speech segmentation depends on words in the listener’s lexicon. For example, word recognition under TRACE assumes that speech will be segmented into words rather than combinations of words and non-words. However, it does not explain how alternative segmentations leading to valid word combinations are reconciled—for example, distinguishing “Grade A” from “grey day”. This example is problematic for the above accounts, because the two segmentations are phonetically identical, acoustically similar, and are both valid word combinations in English. Early accounts also ignored the problem of converting the acoustic signal into words or phonemes. Specifically, they assume that phonetic features (McClelland and Elman 1986) or acoustic features that underlies perceptual confusions in human listeners (NAM; Shortlist B) have already been successfully extracted from the signal. In short, accounts of inputs that are not continuous acoustic signals cannot explain findings that acoustically identical signals are perceived as different words or phonemes depending on their context (Miller and Liberman 1979, Mann 1980, Holt, Lotto et al. 2000).

Here, we consider speech recognition as a Bayesian inference problem. We introduce a simplified generative model that maps from the continuous acoustic signal (i.e., a time varying auditory signal or spectral fluctuations containing particular formant frequencies) to discrete words using lexical, speaker, and prosodic information. Generating continuous states from a succession of discrete states is a non-trivial issue for a first principle (i.e., ideal Bayesian observer) approach. However, the requisite neuronal message passing can be solved by combining variational (marginal) message passing and predictive coding (a.k.a. Bayesian filtering). This allows one to simulate perception using generative models that entertain mixtures of continuous and discrete states (Friston, Parr et al. 2017, Friston, Rosch et al. 2017).

Previous Bayesian accounts (e.g., Shortlist B: Norris and McQueen 2008) have assumed that listeners use exact Bayesian inference. However, performing the calculations required for exact inference would be difficult for biological systems like ourselves, given the complexity of the speech generation process; see (Friston 2010, Bogacz 2017, Friston, FitzGerald et al. 2017). Appealing to variational inference (Beal 2003) affords a much simpler implementation, which has been applied to a variety of other domains in human perception and cognition (Brown, Friston et al. 2011, Brown, Adams et al. 2013, Parr and Friston 2017). Consequently, speech recognition becomes an optimisation problem that corresponds to minimising variational free energy—or, equivalently, maximising the evidence for a particular generative model.

In this paper, we provide a computational perspective on the segmentation problem—addressing the challenge that there are often several ways in which a sentence can be parsed, and multiple segmentations engender valid word combinations. We therefore treat speech recognition as a problem of selecting the most appropriate segmentation among several alternatives. We assume that the listener selects the segmentation that is least surprising from the perspective of their generative model. In doing so, we cast segmentation as an internal action that selects among competing hypotheses for the most likely causes of the acoustic signal. Although this is a novel computational implementation of speech segmentation, it aligns with the basic idea that competing segmentations are held in working memory before a listener decides on the most appropriate segmentation, as supported by behavioural studies of word recognition in human listeners (Shillcock 1990, Davis, Marslen-Wilson et al. 2002). This idea is similar to that used in previous accounts such as TRACE and Shortlist B. Here, we address the problem of selecting among multiple segmentations of valid word combinations. Our approach accounts for contextual effects using priors; we show that alternative segmentations—such as “Grade A” and “grey day”—can be accounted for by appealing to these (e.g., semantic or contextual) priors.

Conceptualising speech segmentation as an internal (covert) action appeals to the ‘active’ aspect of listening. It is distinct from ‘passive’ listening, which—if truly passive—would not require mental or covert actions. This conceptualisation is grounded in active inference, which has previously been applied to active vision (Grossberg, Roberts et al. 1997, Davison and Murray 2002, Ulanovsky and Moss 2008, Andreopoulos and Tsotsos 2013, Ognibene and Baldassarre 2014, Mirza, Adams et al. 2016, Parr and Friston 2017, Veale, Hafed et al. 2017). Here, we consider the covert placement of word boundaries from the same computational perspective as has been used to model an observer whose task is to decide where to sample the visual scene by making overt saccades (Mirza, Adams et al. 2016, Parr and Friston 2017). The types of computations in this framework therefore appeal to general principles that the brain may use to solve a variety of problems.

This paper comprises four sections, which each describe different elements of active listening. The first section reviews active inference and then describes a simplified but plausible generative model of how (continuous) sound waves are generated from a discrete word with particular (discrete) attributes. The attributes include lexical content, prosody, and speaker characteristics. The division of attributes into lexical, prosodic, and speaker attributes is logical from a generative perspective—and is consistent with neuropsychological studies showing selective deficits in the processing of these attributes (Miller and Liberman 1979, Peretz, Kolinsky et al. 1994). Indeed, these attributes have been considered fundamental characteristics in qualitative models of speech perception such as the ‘auditory face’ model (Belin, Fecteau et al. 2004)—and are known to interact to affect human speech perception (Nygaard, Sommers et al. 1994, Johnsrude, Mackey et al. 2013, Holmes, Domingo et al. 2018). We, therefore, assume these are the types of attributes that human listeners infer when trying to explain the (hidden) causes of an acoustic (speech) signal. This section describes how the generative model can be inverted to determine the most likely lexical, prosodic, and speaker attributes of a word, given a continuous sound wave.

The second section deals with the speech segmentation problem, which becomes important when recognising words within sentences, rather than individual words. It considers the question: how do we determine the most likely onsets and offsets of words within a sentence? For example, how do we parse auditory input to disambiguate “Grade A” from “grey day”? To address this question, we use simple acoustic properties to identify plausible word boundaries. We then appeal to the ‘active’ element of active inference, considering the (implicit) placement of word boundaries as a covert ‘action’. This allows us to use established inference schemes to select among competing segmentations (i.e., hypotheses about different word boundaries). These inference schemes essentially ask: which of the possible segmentations minimise free energy or, equivalently, provide the greatest evidence for the listener’s (internal) model of how words are generated? It is at this point that the relationship between the generative model from the first section and ‘active’ speech segmentation becomes clear: these different elements work in unison when inferring words within a sentence. The generative model operates at the individual word level, whereas speech segmentation operates at the sentence level: the best speech segmentation will maximise the combined evidence for attributes of constituent words. This section concludes with an illustration of the face validity of the active listening scheme by comparing speech recognition (i.e., lexical inference) with and without prior beliefs about the sequence of plausible words that could be encountered—demonstrating how different segmentations that contain valid English words can be disambiguated.

The third section highlights an aspect of speech recognition that has not been simulated under previous accounts. We show that a quantity within active listening can predict neurophysiological responses of the sort measured by electromagnetic recordings (Hasson, Yang et al.) or functional magnetic resonance imaging (fMRI). In particular, the magnitude of belief updating in active listening appears to capture the fluctuations in evoked (or induced) responses that have been demonstrated empirically; e.g., the mismatch negativity (Garrido, Kilner et al. 2009, Morlet and Fischer 2014), P300 (Donchin and Coles 1988, Morlet and Fischer 2014), and N400 (Kutas and Hillyard 1980). Broadly speaking, this suggests that elements of speech perception are consistent with predictive coding (see (Poeppel and Monahan 2011) for a review). Formally, belief updating is related to the difference between *prior* beliefs about states in the generative model to *posterior* beliefs. In other words, the amount that beliefs change after sampling sensory evidence. This is variously known as *Bayesian surprise*, salience, information gain, or complexity. In this section, we illustrate the similarity between belief updates and violation responses, showing that the magnitude of belief updating depends upon prior expectations about particular words in the lexicon (Cole, Jakimik et al. 1980, Mattys and Melhorn 2007, Mattys, Melhorn et al. 2007, Kim, Stephens et al. 2012) and the quality of sensory evidence; e.g., when speech is acoustically masked by background noise (“speech-in-noise”) (Sams, Paavilainen et al. 1985, Winkler, Denham et al. 2009). We conclude by discussing how the model could be developed for future applications, and its potential utility in the cognitive neuroscience (and neuropsychology) of auditory perception and language.

## A generative model of spoken words

Active inference is a first principle account of action and perception in sentient creatures (Friston, FitzGerald et al. 2017). It is based upon the idea that synaptic activity, efficacy and connectivity all change to maximise the evidence for a model of how our sensations are generated. Formally, this means treating neuronal dynamics as a *gradient flow* on a quantity that is always greater than (negative) log evidence (Friston, Parr et al. 2017). This quantity is known as variational free energy in physics and statistics (Feynman 1972, Hinton and Zemel 1993). The complement (i.e., negative) of this quantity is known as an evidence lower bound (ELBO) in machine learning (Winn and Bishop 2005). A gradient flow is simply a way of writing down dynamics in terms of equations of motion that ensure a certain function is minimised— in this case, variational free energy. The resulting dynamics furnish a model of neuronal fluctuations (and changes in synaptic efficacy and connectivity) that necessarily minimise free energy or maximise model evidence. In short, if one simulates speech recognition using active inference, one automatically provides an account of the accompanying neuronal dynamics.

This approach to understanding and modelling (active) inference in the brain has been applied in many settings, using exactly the same schemes and principles. The only thing that distinguishes one application from another is the form of the generative model. In other words, if one can write down a probabilistic model of how some sensory input was generated, one can invert the model—using standard model inversion schemes—to simulate neuronal dynamics and implicit belief updating in the brain: See (Friston, Parr et al. 2017) for a detailed summary of these schemes that cover models of both discrete and continuous states generating sensations. See also (Bastos, Usrey et al. 2012, Friston, FitzGerald et al. 2017) for a discussion of neurobiological implementation, in terms of attending process theories, for continuous and discrete state space models, respectively.

In this section, we focus on the form of a (simplified) generative model that can be used to generate continuous acoustic signals associated with a particular word. A benefit of this active inference approach is that the generative model can be used to both generate synthetic speech (by applying the forward model) and recognise speech (by inverting the model). The goal is not to provide a state-of-the art speech synthesis system, but rather to use the generative model and accompanying inference scheme to simulate listening behaviour and neural responses. The work reported in this paper is a prelude to a model of natural language processing, in which the current generative model is equipped with higher levels to enable dyadic exchanges; namely, conversations that entail questions and answers that resolve uncertainty about shared narratives or beliefs. In this paper, we restrict ourselves to inference about sequences of words—and assume that simulated subjects are equipped with prior beliefs about which words are more or less likely in a short sentence or phrase. In a more complex (i.e., deep hierarchical) model, these beliefs would be available from a higher level. These prior beliefs are about the likely semantic content of spoken words; for example, based on previous words in a sentence (Dubno, Ahlstrom et al. 2000) or the topic of conversation (Holmes, Folkeard et al. 2018). Note that previous accounts of speech recognition, such as Shortlist B (Norris and McQueen 2008), assume that priors reflect only word frequency, rather than priors that can be flexibly updated based on context. Technically, these kinds of context-sensitive priors are known as empirical priors—and are an integral part of hierarchical generative models.

In this paper, we deal with the lowest level of the generative model; namely, given a particular lexical content, prosody and speaker identity, how would one generate a spoken word in terms of its acoustic timeseries. In the next section of this paper, we turn to the problem of segmentation (i.e., identifying word boundaries) and the enactive aspects of the current scheme. It will become apparent later on that these two (perceptual and enactive) aspects of active listening go hand-in-hand.

Figure 1 summarises the modelling of a spoken word, from the perspectives of generation and recognition. The model considers: how is an acoustic signal generated given the causes of a spoken word, in terms of ‘what’ word is spoken (*lexical*), ‘how’ it is spoken (*prosody*), and ‘who’ is speaking (*speaker* identity)? From the perspective of word generation, it takes *lexical*, *speaker*, and *prosody* parameters and generates an expected acoustic signal. The *lexical* state consists of frequency and temporal coefficients corresponding to words in the lexicon. The model includes two *speaker* states: fundamental frequency and formant scaling. It includes four *prosody* states: amplitude, duration, timbre, and inflection. Within each of these states, different factors correspond to different lexical items, or the fundamental frequency associated with different speakers, for example.

**Figure 1.**
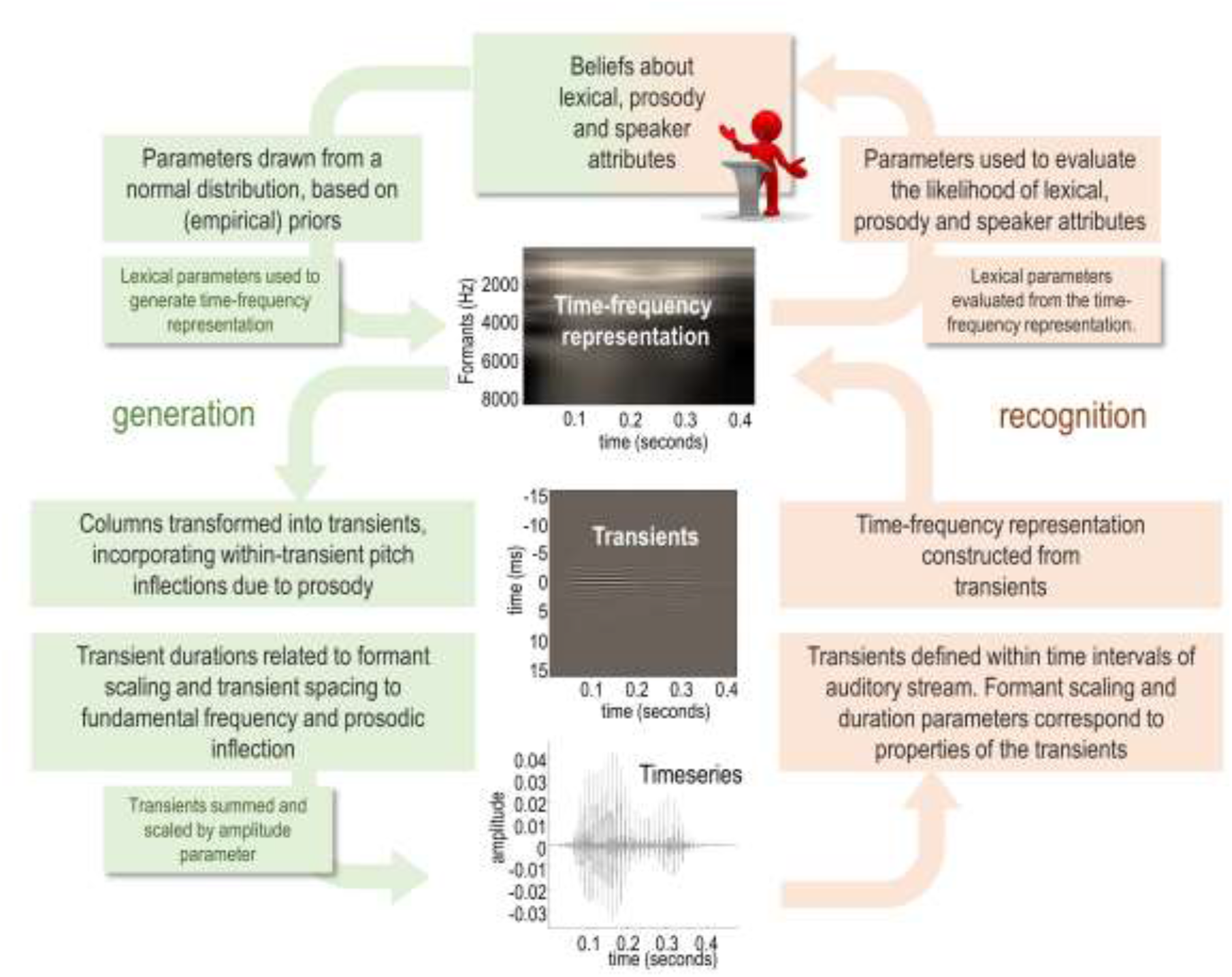
A generative model of a word. This figure illustrates the generative model from the perspective of word generation (green panels) and accompanying inversion (orange panels), which corresponds to word recognition. For the equations describing these probabilistic transformations, please see Appendix 1.

The model starts by sampling parameters from a set of probability distributions, which are modelled as separate Gaussians. The means and covariances of the Gaussians have been specified in advance; they can be entered into the model explicitly (by hand) or they can be estimated empirically based on training samples of speech. Sampling parameters from distributions with particular means and variances accounts for the fact that the same lexical item spoken by the same speaker with the same prosody does not always produce an identical acoustic signal, and—conversely—because the distributions are allowed to overlap, a similar acoustic signal can be generated by different combinations of factors. The (discrete) lexical content of a word is sampled from a (categorical) probability distribution over words in a lexicon. This is based on how likely particular words are to be spoken. Ultimately, the selected parameters are combined, in a nonlinear way, to generate an acoustic timeseries corresponding to the articulated word.

The acoustic timeseries is generated from a sequence of transients, whose properties are determined by the selected parameters. Each word (i.e., *lexical* item) is associated with a matrix of frequency and temporal coefficients (for a discrete cosine transform) that can be used to generate a time-frequency representation of the spoken word (i.e., the spectrogram) when combined with *speaker* and *prosody* information. Each column of the time-frequency representation is used to generate a transient. These transients can be thought of as pulses or ‘shockwaves’ at the glottal pulse rate, which are modulated by the shape of the vocal tract. The instantaneous fundamental frequency is related to the average fundamental frequency of a particular speaker, but also varies smoothly over time based on inflections due to prosody. The prosodic inflection parameters encode: (1) the average fundamental frequency relative to the speaker average, (2) increases or decreases in fundamental frequency over time, and (3) the acceleration or deceleration of changes in fundamental frequency. The instantaneous fundamental frequency determines the spacing of the transients. The durations of the transients are determined by the formant frequencies, which depend on the lexical parameters and the speaker formant scaling parameter. The formant frequencies correspond to the frequency bins in the time-frequency representation. The number of transients that are aggregated to construct the timeseries is determined by the time intervals in the time-frequency representation. Figure 2 provides an illustration of how a sequence of transients is generated. In the final step, the transients are summed together and scaled by an amplitude parameter. For mathematical detail, the equations corresponding to the generative model are shown in Figure 11 and are described in Appendix 1. For an algorithmic description, please see the demonstration (annotated Matlab) code—that reproduces the simulations below—which can be read as pseudocode (see Software note).

**Figure 2.**
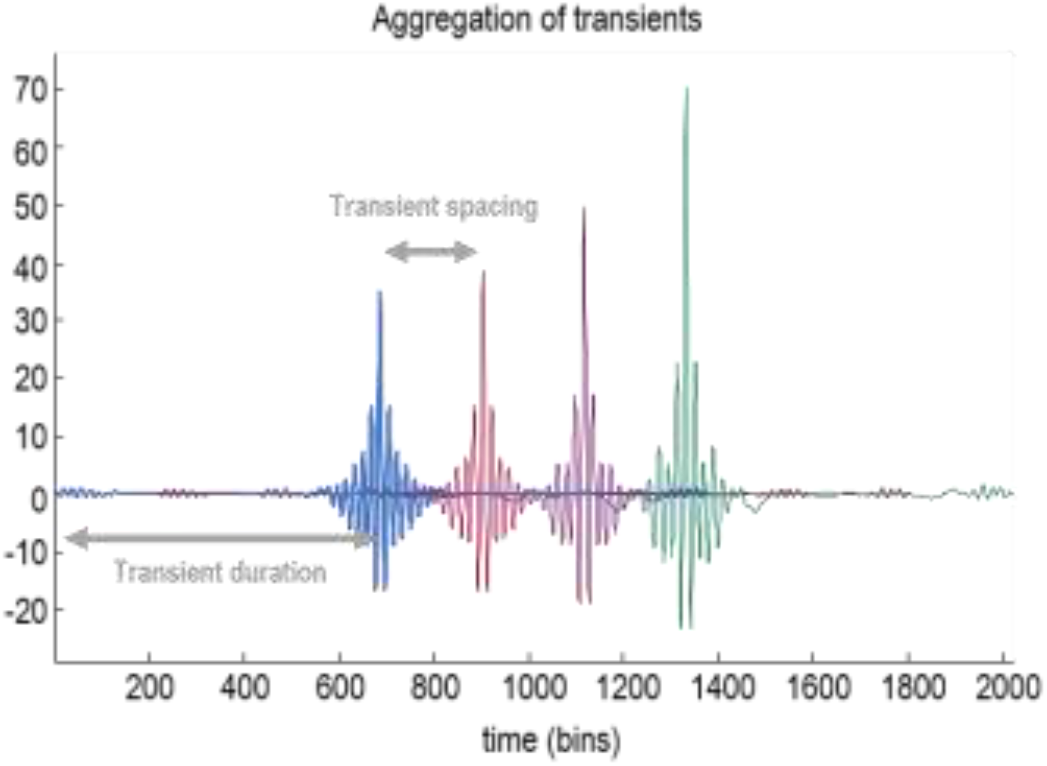
Fundamental and formant intervals. This figure illustrates the way in which an acoustic timeseries is generated by assembling a succession of transients separated by an interval that is inversely proportional to the (instantaneous) fundamental frequency. The duration of each transient places an upper bound on the wavelength of the formant frequencies—and corresponds to the minimum frequency, which we take to be the first formant frequency.

In effect, the lexical parameters—which, under this generative model, determine the formant frequencies— parameterise a trajectory through high-dimensional formant frequency space, which becomes apparent as the word unfolds. The prosody of the word determines the duration and inflection of the fundamental interval function, while speaker identity determines the average fundamental frequency—which relates to the interval between transients—and a formant scaling parameter that determines the duration of each transient. With such a model in place, one can, in principle, generate any word, spoken with any prosody by any speaker, by sampling the correct parameters from their appropriate distributions. In what follows, we briefly review the inversion of this model given an acoustic timeseries.

### Model inversion or word recognition

Now we have established a generative model that is capable of producing a spoken word, word recognition can be achieved by inverting the model. This section describes a plausible inversion scheme in the context of our particular generative model of spoken words. In principle, given any generative model it should be possible to use Bayesian model inversion to invert the timeseries, using generalised (variational or Bayesian) filtering; also known as predictive coding (Norris, McQueen et al. 2016). However, given we have assumed a deterministic generation of acoustic signals from parameters, we know that the posterior beliefs about parameters will take the form of Dirac delta functions, whose only parameter is a mode. This means that in practice, it is simpler to cache an epoch of the timeseries and use *maximum a posteriori* (Kim, Frisina et al.) estimates of the parameters, based upon least squares. One can then evaluate the posterior probability of discrete *lexical*, *prosody* and *speaker* states, using the respective likelihood of the (Kim, Frisina et al.) parameter estimates (and any priors over discrete states should they be available). This MAP scheme can be read in the spirit of predictive coding that has been *amortised* (Zhang, Butepage et al. 2018). In other words, the inversion scheme reduces to a nonlinear recognition function—a series of equations that map from epochs of the acoustic signal to parameters encoding lexical content, prosody and identity.

Model inversion rests on the assumption that we have isolated the acoustic timeseries corresponding to an individual word. The next section deals with the segmentation problem, which involves enactive processes. For now, we will assume that we have identified an epoch of the acoustic signal that might plausibly contain one word—and that we wish to evaluate the probabilities of *lexical*, *prosody*, and *speaker* states within this epoch.

In brief, the recognition scheme comprises the following steps (see Figure 1). The instantaneous frequency is estimated by first calculating ‘fundamental intervals’, which are the reciprocal of the instantaneous frequency. The fundamental intervals are calculated by bandpass filtering the acoustic signal around the prior value for the speaker fundamental frequency parameter; the positions of peaks in the filtered signal correspond to the fundamental intervals. Please see Figure 3 for an illustration of how the fundamental intervals are estimated and Figure 4 to see the fundamental frequency and formant frequencies projected onto the spectrum of a speech sample.

**Figure 3.**
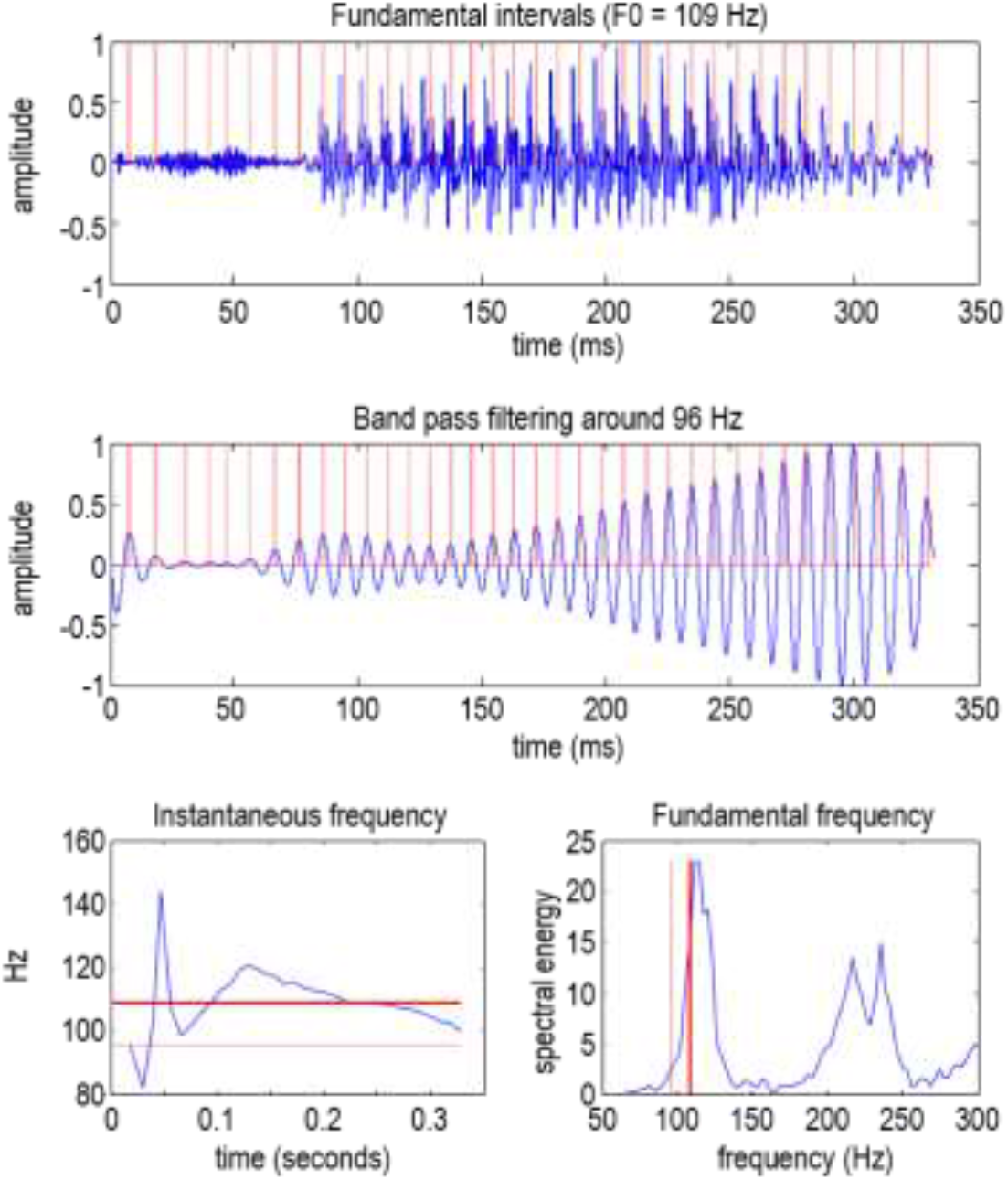
Fundamental frequencies and intervals. This figure illustrates the estimation of fluctuations around the fundamental frequency during the articulation of (the first part of) a word. These fluctuations correspond to changes in the fundamental interval; namely, the reciprocal of the instantaneous frequency. The upper panel shows the original timeseries, while the middle panel shows the same timeseries after bandpass filtering. The peaks (i.e., phase crossings) then determine the intervals, which are plotted in terms of instantaneous frequencies on the lower left (as a blue line). The solid red line corresponds to the mean frequency (here, 109 Hz), while the broken red line corresponds to the centre frequency of the bandpass filtering (here, 96 Hz) which is centred on the prior for the speaker average fundamental frequency. The same frequencies are shown on the lower right panel, superimposed on the spectral energy (the absolute values of the accompanying Fourier coefficients of the timeseries in the upper panel). The ensuing fundamental intervals are depicted as red lines in the upper two panels.

**Figure 4.**
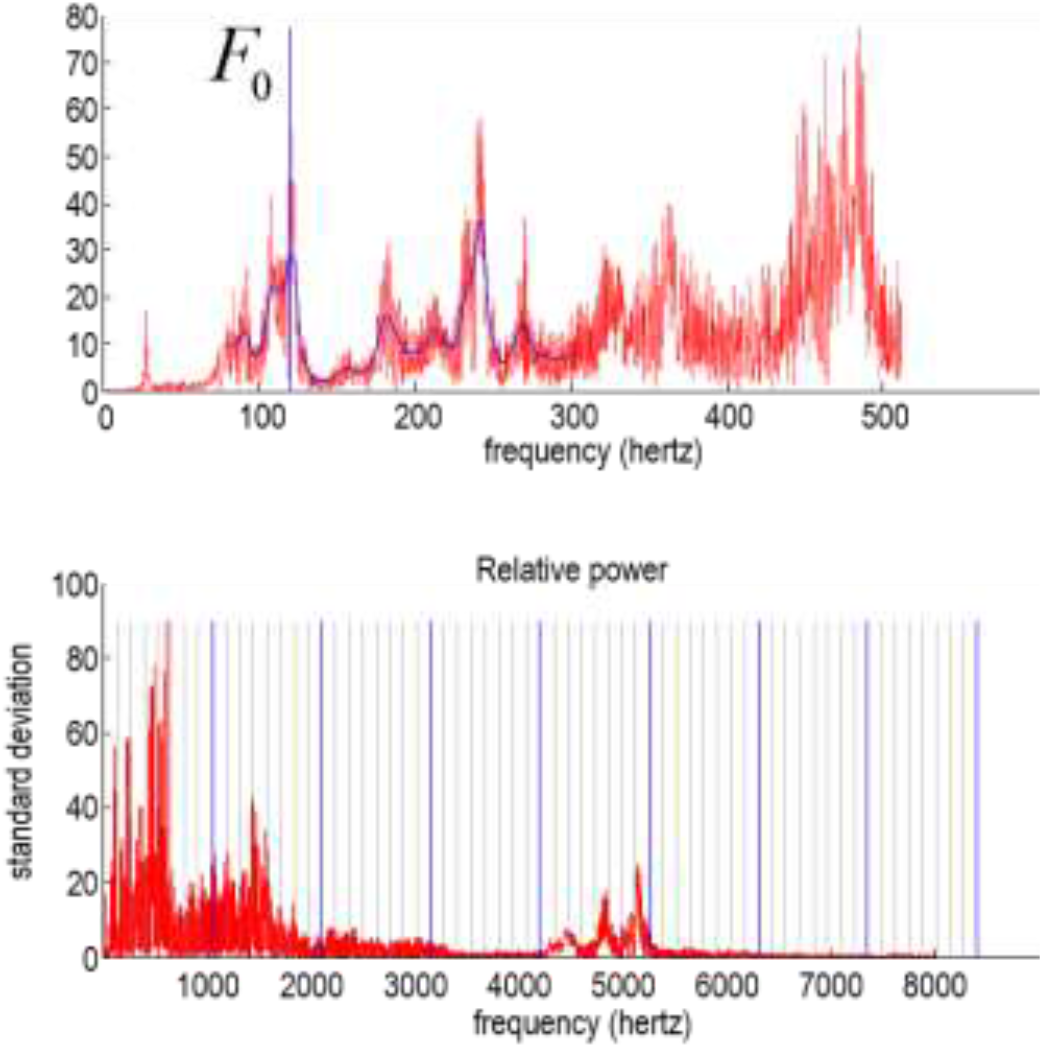
Fundamental and formant frequencies: Both plots show the root mean square power (i.e., absolute value of Fourier coefficients) following the Fourier transform of a short segment of speech. The frequency range in the upper plot covers the first 500 Hz. The first peak in power (illustrated by the blue vertical line) corresponds to the *fundamental frequency*, which is typically between 80 and 150 Hz for adult men and up to 350 Hz for adult women. The lower panel shows the same spectral decomposition but covers 8000 Hz to illustrate formant frequencies. The solid blue lines show the calculated formant frequency and its multiples, while the grey lines arbitrarily divide the frequency intervals into eight bins. These frequencies define the frequencies used for the spectral decomposition.

Next, the inversion scheme essentially deconstructs transients (i.e., segments) from the epoch. The formant frequencies are estimated by evaluating the cross-covariance function over short segments; the length of the segments is the inverse of the first formant frequency and the segments are centred on each fundamental interval. This is based on the simplifying assumption that the spectral content of each transient, within each segment, is sufficient to generate the word. The formant frequencies are then used to project back to a time-frequency representation.

To infer the lexical content, prosody and speaker, the parameter estimates from the nonlinear transformations above can be used to evaluate the likelihood of each discrete attribute. This likelihood is then combined with a prior to produce a posterior categorical distribution over the attributes in question. For the *lexical* content of the word, this just corresponds to an index in the lexicon. Here, the lexicon is assumed to be small for simplicity, although it would be trivial to extend the model to accommodate more comprehensive lexicons. The likelihood is based upon the mean and precision (i.e., inverse covariance) of the lexical parameters in the usual way, where the sufficient statistics of this (likelihood) model—for each word—are evaluated using some exemplar or training set of words. This completes the description of word recognition based upon the generative model above. For details of the equations used in model inversion, please see Appendix 2.

In summary, the above transformations simply reverse the operations used for word generation in the previous section. The combination of prior expectations with the likelihoods of each attribute is a key feature of this inversion scheme that will allow the model to accommodate contextual effects on speech recognition. In other words, we are more likely to interpret speech consistent with our prior expectations. This will become evident in the simulations later in this paper.

After the discrete parameters have been inferred from a continuous timeseries through model inversion, they could be entered back into the generative model to synthesise a new timeseries that would share some properties with the timeseries that was used to infer the discrete parameters. This simply involves projecting the lexical coefficients back into a time frequency representation, implementing the inverse discrete cosine transform to produce (after scaling with the timbre parameter and exponentiation) a series of (time symmetric) transients, which are aggregated to form the acoustic timeseries. This is essentially what is illustrated in Figure 1. Indeed, the processes of inversion and generation can be iterated (see below) to check the fidelity of the forward and inverse transformations that map between the acoustic timeseries and formant representation.

## Speech segmentation as an active process

So far, we have a generative model (and amortised elements of a predictive coding scheme) that generates an appropriate time series, given discrete *lexical* (i.e., what), *prosody* (i.e., how) and *speaker* (i.e., who) states (i.e., latent causes of the word). It can also be inverted to infer the attributes of a word given an acoustic timeseries. However, in our everyday lives, we usually hear series of words rather than words in isolation. In this section, we combine the generative model with an active segmentation process, to infer the most likely *sequence* of words given a continuous timeseries.

This requires us to address the following problem: we have not specified how the onsets and offsets of the interval containing the word are generated (i.e., when). Clearly, there are some prior constraints on the generation of these intervals. For example, the offset of one word should precede the onset of the subsequent word. Furthermore, the intervals contained between the onset and offset must lie in some plausible time range. We also know that segmentations are more likely to contain words than non-words (Ganong 1980, Billig, Davis et al. 2013), and listeners have prior knowledge of the words that are possible in a language (‘possible word constraint’) (Norris, McQueen et al. 1997). In the current segmentations, we account for these simple constraints and, effectively, offload inference about word boundaries to the *active* part of active inference. The only acoustic cue we use is the contour of the amplitude envelope, which has previously been identified as a cue that human listeners use for speech segmentation (Lehiste 1960).

In brief, we assume that boundary segmentations are not entirely specified by the acoustic signal, and conceptualise the segmentation problem as a problem of choosing which boundaries to select given several possible segmentations; in a similar way as we would select visual actions (e.g., saccadic eye movements or oculomotor pursuit) to fixate or track a visual object given multiple possible actions. In the current setting, this simply means identifying a number of plausible boundary intervals and finding the interval that provides the greatest evidence for our prior beliefs about the words we hear. This is the same principle used to explain motor and autonomic action under active inference (Friston, Mattout et al. 2011). For example, classical motor reflexes can be construed as minimising proprioceptive prediction error (i.e. minimising variational free energy or maximising model evidence) as described in (Adams, Shipp et al. 2013). Formally identical arguments have been applied in the setting of interoceptive inference where motor reflexes are replaced by autonomic reflexes that realise autonomic set-points or homoeostasis (Seth 2014).

In the current context, we essentially treat the decision about speech segmentation as a covert action from a computational perspective, which shares similarities with the overt actions used in other settings. This can be implemented in a straightforward fashion by selecting boundary pairs (i.e., offsets and onsets) and evaluating their free energy under some prior beliefs about the next word. Ultimately, we want to select the boundary pairs with the smallest free energy—which effectively selects the interval with the greatest evidence (a.k.a., marginal likelihood) of auditory outcomes contained in that interval. This follows because the variational free energy, by construction, represents an upper bound on log evidence (see Appendix 3 for more details and the corresponding equations). Importantly, both posterior beliefs about latent states (i.e., *lexical*, *prosody*, and *speaker*) and the active selection of acoustic intervals optimise free energy. This is the signature of active inference. In this instance, the posterior beliefs obtain from the likelihood of the lexical, prosody and identity parameters, given the associated states.

For words spoken in isolation, one can identify candidate boundaries using threshold crossings of the amplitude envelope (where the threshold is a low value, roughly corresponding to the noise floor). However, it is well known that a continuous stream of words does not always contain ‘silent’ (i.e., below-threshold) gaps between words and, conversely, silence can occur between two syllables of the same word. We therefore include local minima of the amplitude envelope as candidate boundaries. It is important to note that these are only *candidate* boundaries—in other words, plausible hypotheses for segmentations of the acoustic signal. We will turn to the question of which interval is *selected* later, during which candidate segmentations are combined with (lexical) priors. In practice, this means that two syllables separated by a silent gap are not always classified as separate words—consistent with the knowledge that naturally spoken words often contain silent gaps that—to a naïve listener—could be confused with word boundaries. An example of the candidate boundary points is illustrated in Figure 5. Please see figure legend for details.

**Figure 5.**
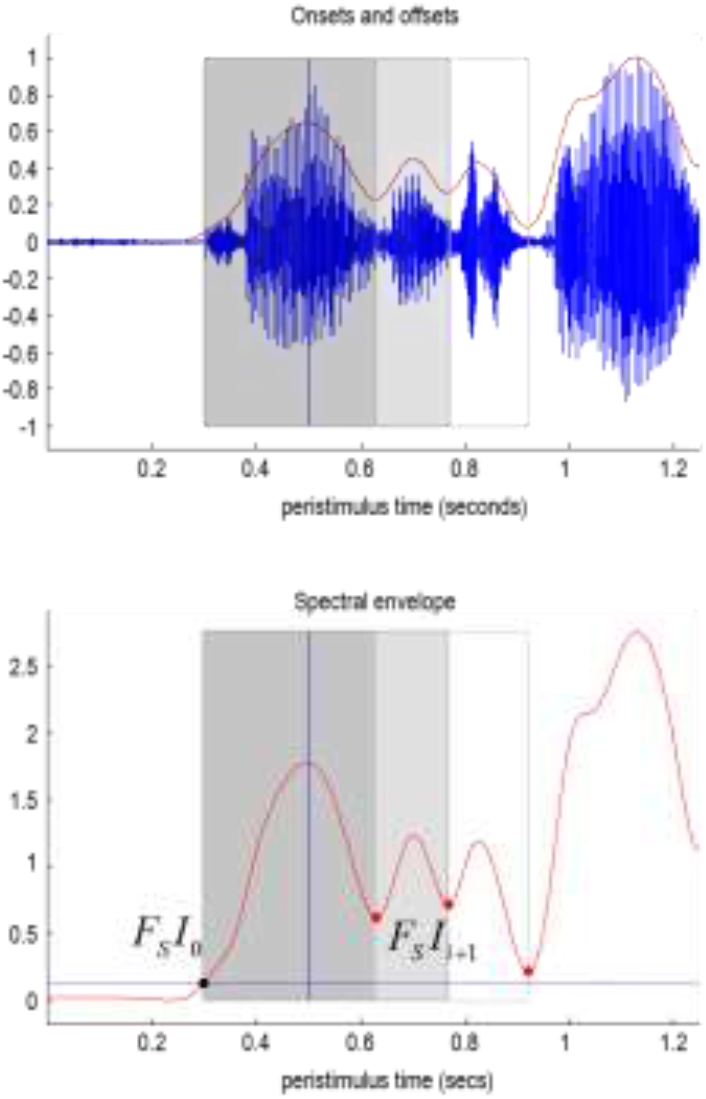
Spectral envelopes and segment boundaries. This figure provides an example of how candidate intervals containing words are identified using the spectral envelope. The upper panel shows a timeseries produced by saying “triangle, square”. The timeseries is high pass filtered and smoothed using a Gaussian kernel. The dotted red line in the upper panel shows the resulting spectral envelope, after subtracting the minimum. The broken line corresponds to a threshold: 1/16^th^ of the maximum encountered during the (1250 ms) epoch. This envelope is reproduced in the lower panel (red line). Boundaries are then identified as the first crossing (black dot) of the threshold (horizontal blue line) before the spectral peak and the last crossing after the peak. These boundaries are then supplemented with the internal minima between the peak and offset (red dots). These boundaries then generate a set of intervals for subsequent selection during the recognition or inference process. Here, there are three such intervals. The first contains the first two syllables of triangle, the second contains the word “triangle”. The third additionally includes the first phoneme of “square”. In this example, the second interval was selected as the most plausible (i.e., free energy reducing) candidate to correctly infer that this segment contained the word “triangle”. The vertical blue line corresponds to the first spectral peak following the offset of the last word, which provides a lower bound on the onset.

Using this procedure to identify candidate intervals, one can select the interval that minimises free energy (or has the greatest evidence under prior beliefs about the next word). In other words, for each candidate interval, the likelihood of the lexical parameters is evaluated—for all plausible words—to create a belief over lexical content, in terms of a probability distribution. This posterior belief is then used to evaluate the log evidence (i.e., free energy) of each interval. The interval (and associated posterior beliefs) with the greatest evidence is selected. The offset of this interval specifies the onset of the next segment and the process starts again.

Treating speech segmentation as a problem of (covertly) sampling among plausible intervals is interesting from a mathematical perspective. The free energy associated with a particular action is a trade-off between the accuracy of sensory observations under the generative model and the complexity of belief updating on the basis of those observations (see Appendix 3 for the equations). In the current setting, these quantities can be evaluated explicitly, because the evidence has already been accumulated. Thus, the accuracy term simply scores the expected log likelihood of the auditory observations under posterior beliefs about the lexical categories that generated them. The complexity term scores the difference between the prior beliefs and the new beliefs based on auditory observations. This will become an important quantity later and, essentially, reflects the degree of belief updating associated with selecting one lexical parsing over another. Phrased another way, the goal of segmentation under active listening is to sample data in a way that requires the most parsimonious degree of belief updating, in accord with Ockham’s principle (Maisto, Donnarumma et al. 2015).

Figure 6 shows the consequence of this form of active listening by comparing segmentation and recognition with and without appropriate prior beliefs (please see the figure legend for details). The input to this simulation is a continuous acoustic signal that has alternative parsings, leading to different lexical segmentations. The timeseries in Figures 6A and 6E are identical, but the segmentation (as indicated by the colours) differs. The point of this simulation is to show that the selected segmentation depends on the distribution of the priors. When the artificial listener has no particular prior beliefs about which words will be heard (left panel), the priors are uniform, and recognition goes awry after the first two words (“triangle square”). The scheme inferred that the best possible explanation for the subsequent words was a series of shorter words (“a is red a is red”; Figure 6B). From Figure 6C, we can tell that the artificial listener was uncertain about the correct parsing—reflecting the fact that this signal was difficult to segment because there were several parsings that would be plausible in English (displayed as grey shaded regions). However, when the artificial listener was equipped with strong prior beliefs that the words they would hear would be shape words (the words “triangle” and “square”), it recovered the correct parsing (“triangle square triangle square triangle square”; Figure 6F). Note that the acoustic boundaries for these two lexical segmentations differ—highlighting that speech segmentation and lexical inference go hand-in-hand, under this framework.

**Figure 6.**
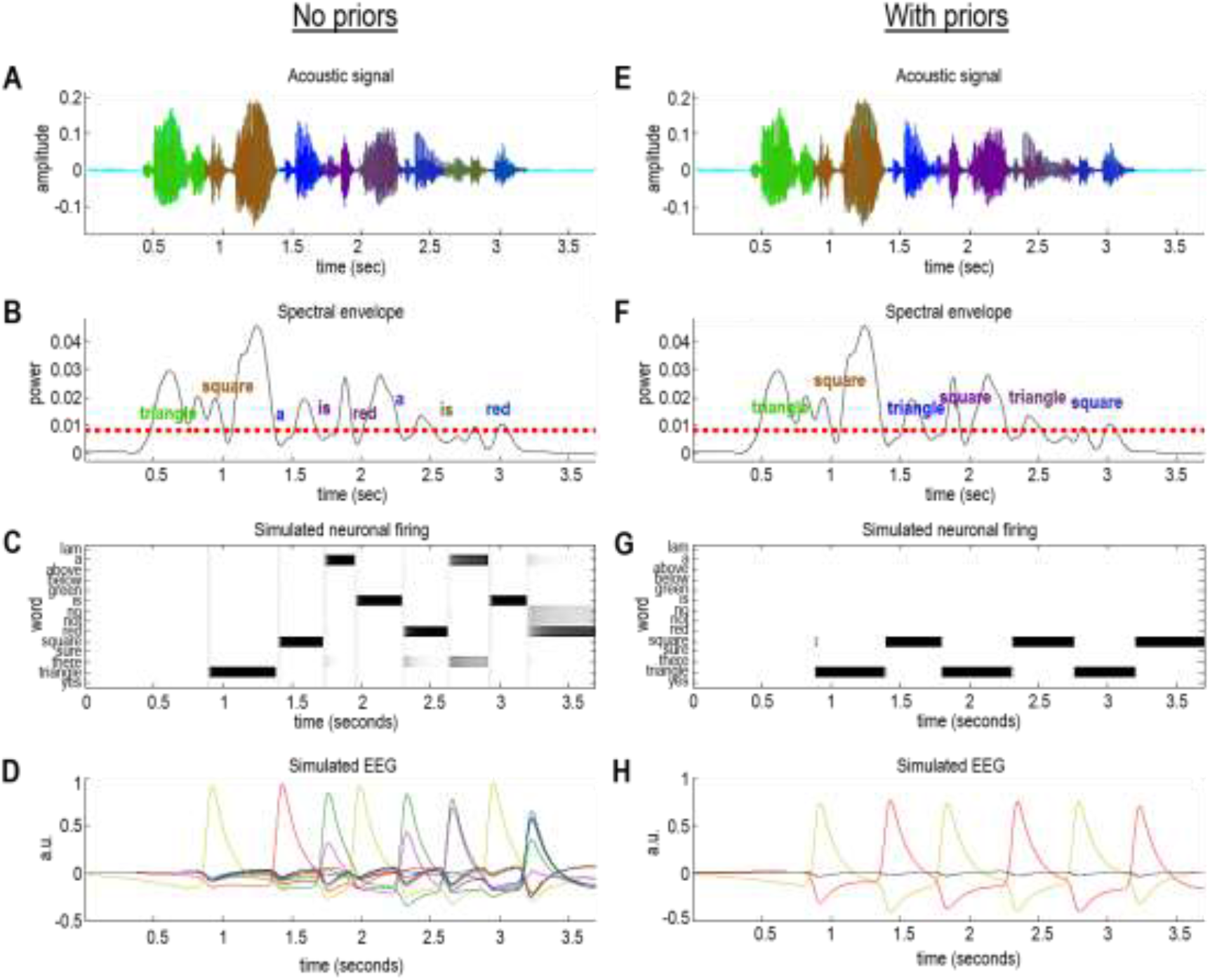
Speech recognition and segmentation. **Left panel**: This panel shows the results of active listening to a sequence of words: a succession of “triangle, square, triangle, square….”. Its format will be used in subsequent figures and is described in detail here. Panel A shows the acoustic timeseries as a function of time in seconds. The different colours correspond to the segmentation selected by the active listening scheme, with each colour corresponding to an inferred word. Regions of cyan denote parts of the timeseries that were not contained within a word boundary. Panel B shows the accompanying spectral envelope (back line) and the threshold (red dashed line) used to identify subsequent peaks. The first peak of each successive word centres the boundary identification scheme of Panel A. The words that have been inferred are shown in the same colours as the upper panel at their (inferred) onset. Panels C–D show the results of simulated neuronal firing patterns and local field potentials or electroencephalographic responses. These are based upon a simple form of belief updating cast as a neuronally plausible gradient descent on variational free energy (please see main text). Panel C shows the activity of neuronal populations encoding each potential word (here, 14 alternatives listed on the Y axis). These are portrayed as starting at the offset of each word. Effectively, these reflect a competition between lexical representations that record the selection of the most likely explanation. Sometimes this selection is definitive: for example, the first word (“triangle”) supervenes almost immediately. Conversely, some words induce a belief updating that is more uncertain. For example, the last word (“red”) has at least three competing explanations (i.e., “no”, “not” and “a”). Even after convergence to a particular posterior belief, there is still some residual uncertainty about whether “red” was heard. Note that the amplitude of the spectral envelope is only just above threshold. In other words, this word was spoken rather softly. Panel D shows the same data after taking the temporal derivative and filtering between 1 and 16 Hz. This reveals fluctuations in (simulated) depolarisation that drives the increases or decreases in neuronal firing of the panels above. In this example, the sequence of words was falsely inferred to be a mixture of several words not actually spoken. This failure to recognise the words reflects the fact that the sequence was difficult to parse or segment. Once segmentation fails, it is difficult to pick up the correct sequence of segmentations that will, in turn, support veridical inference. These results can be compared with the equivalent results when appropriate priors are supplied to enable a more veridical segmentation and subsequent recognition. **Right panel**: This panel shows the results of active listening using the same auditory stream as in the left panel. The only difference here is that the (synthetic) subject was equipped with strong prior beliefs that the only words in play were either “triangle” or “square”. This meant that the agent could properly identify the succession of words, by selecting the veridical word boundaries and, by implication, the boundaries of subsequent words. If one compares the ensuing segmentation with corresponding segmentation in the absence of informative priors, one can see clearly where segmentation failed in the previous example. For example, the last word (i.e., “square”) is correctly identified in dark blue in Panel F. Whereas, in Panel B (without prior constraints), the last phoneme of the word “square” was inferred as “red” and the first phoneme was assigned to a different word (“is”). The comparative analysis of these segmentations highlights the ‘handshake’ between inferring the boundaries in a spectral envelope and correctly inferring the lexical content on the basis of fluctuations in formant frequencies.

These two examples are analogous to the “Grade A” versus “grey day” example that we considered in the introduction. As in our simulated example, there is no consistent acoustic cue that differentiates “Grade A” from “grey day”—and, therefore, priors play an essential disambiguating role. The active segmentation would identify these two (and perhaps additional) possible segmentations, and the percept would be the one that was most similar to the priors. In other words, these two segmentations would be distinguished by different prior beliefs, which could originate from a higher (semantic or contextual) level—for example, whether the topic of conversation was about the weather or a student’s exam results. In a comprehensive treatment, these would be empirical prior beliefs generated by deep temporal models of the sort described in (Kiebel, Daunizeau et al. 2009, Friston, Rosch et al. 2017). For simplicity and focus, we assume here that priors about sequential lexical content—of the sort that could be formed by lexical and semantic predictions—are available to a subject in the form of categorical probability distributions.

### Belief updating and neuronal dynamics

Figure 6 includes a characterisation of simulated word recognition in terms of neuronal responses (Figure 6C–D, G–H). These (simulated) neuronal responses inherit from the neuronal (marginal) message passing scheme described in (Friston, Parr et al. 2017, Parr, Markovic et al. 2019). They reflect belief updating about the lexical category for each word; the simulated neuronal responses are simply the gradient flow on free energy that is associated with belief updating in active listening. The prediction error is the (negative) free energy gradient that drives neuronal dynamics. Mathematically, the prediction error is the difference between the optimal log posterior and current estimate of this. As detailed in Appendix 3, log expectations about hidden states can be associated with depolarisation of neurons or neuronal populations encoding expectations about hidden states, while firing rates encode expectations *per se*.

Figure 6 reproduces these simulated neuronal responses following the processing of each word. These responses are shown in terms of spike rates, as would be recorded with single unit electrodes (Figure 6C, G) and depolarisation that would be measured with EEG (Figure 6D, H). Under this formulation, neuronal activity starts off from some prior expectations and evolves, via a gradient flow on free energy (i.e., prediction error) to encode posterior expectations. Because depolarisation corresponds to the rate of change of these beliefs (expressed as log expectations) they show peak responses during the greatest degree of belief updating from priors to posterior expectations. After filtering, the simulated depolarisations look like evoked responses that are typically observed in human studies (as discussed in more detail below).

### Summary

The message from the simulations in Figure 6 is that proper segmentation and subsequent inference about lexical content obtain only with particular priors. If we remove prior constraints entirely, the synthetic listener failed to identify the correct intervals; it falsely inferred the presence of words that were not uttered and ‘missed’ words that were spoken. It is worth mentioning that the absence of priors would be extremely unlikely in realistic contexts, because our knowledge of language generates expectations about plausible words in any given sentence (e.g., due to syntactic and semantic constraints, as well as simple effects of word frequency) and contextual knowledge (e.g., knowing the topic of conversation, or being in a particular setting) will also supply empirical priors. Indeed, the effect of priors on speech segmentation is well-established in human speech perception. The common observation that word boundaries are difficult to ascertain in an unknown language is an intuitive example that priors based on lexical knowledge help to determine speech segmentation. In addition, the way that humans segment speech depends on previous words in a sentence (Cole, Jakimik et al. 1980, Mattys and Melhorn 2007, Mattys, Melhorn et al. 2007, Kim, Stephens et al. 2012)—a simple demonstration that priors are flexibly applied in different contexts. The aim of this simulation was to demonstrate the role of priors in speech recognition under active listening.

This simulation also shows that active listening goes beyond simply inferring the best explanation for a particular sensory signal: active listening also infers which signals to ‘sample’. By this, we mean that different segments (corresponding to plausible word boundaries) of the speech signal are evaluated, with the goal of ‘sampling’ or selecting one set of intervals. The action (here, covert placement of word boundaries, which can be considered more generally as active sampling) therefore goes hand-in-hand with perception. This is demonstrated in the left panel of Figure 6: Although the words recognised provide the best (Kim, Frisina et al.) explanation for acoustic sensations, both the words themselves and the placement of word boundaries are categorically different from the right panel of Figure 6, in which the model was equipped with different (uniform) prior beliefs. This ability to integrate different levels of beliefs and inference is consistent with a hierarchical architecture, as suggested by (i) experimental studies that have measured brain responses during speech perception (Davis and Johnsrude 2003, Vinckier, Dehaene et al. 2007, DeWitt and Rauschecker 2012), (ii) studies that examine the weights participants assign to different cue types during speech segmentation; e.g., (Mattys, White et al. 2005), and (iii) cognitive accounts of speech processing (McClelland and Elman 1986, Gaskell and Marslen-Wilson 1997). In the next section, we turn to the electrophysiological correlates of this belief updating and ask what predictions this model of auditory inference can offer.

### Face validity: Simulating sentence recognition

Here, we use the generative model and inversion scheme described above, under simple prior beliefs about a sentence, to illustrate the circular causality implicit in Bayesian belief updating. In brief, we will examine how prior beliefs underwrite word segmentation and how segmentation changes in the absence of appropriate priors. We then look at how the selected speech segmentation updates subsequent prior beliefs and how the ensuing Bayesian surprise may manifest electrophysiologically. To illustrate the effect of priors, we chose the following sentence: “Is there a square above?” This is a completely arbitrary sentence but is interesting because the formant frequencies in the word “square” have a bimodal (biphone) structure (Bashford, Warren et al. 2008), which means there is a fairly severe segmentation problem at hand. Will a simulated subject segment “square” properly or—as in Figure 6-append the first phone to the previous word? If they do infer the words correctly, how do priors manifest in terms of belief updating?

Figure 7 shows the results of integrating the active inference scheme above with strong (left panels) or uniform (right panels) prior beliefs. In this example, prior beliefs were definitive for the first three words (“is there a”) with more ambiguous prior for the last two words: for the fourth word, the possibilities included “square” and “triangle”. For the final word, the possibilities included “above”, “below” and “there”). These priors were selected because they are lexically congruent and represent a plausible belief that a listener might have about the content of a sentence. Please see the figure legend for technical details. The message from this simulation is that priors play a key role in resolving uncertainty and subsequent competition among neuronal representation.

**Figure 7.**
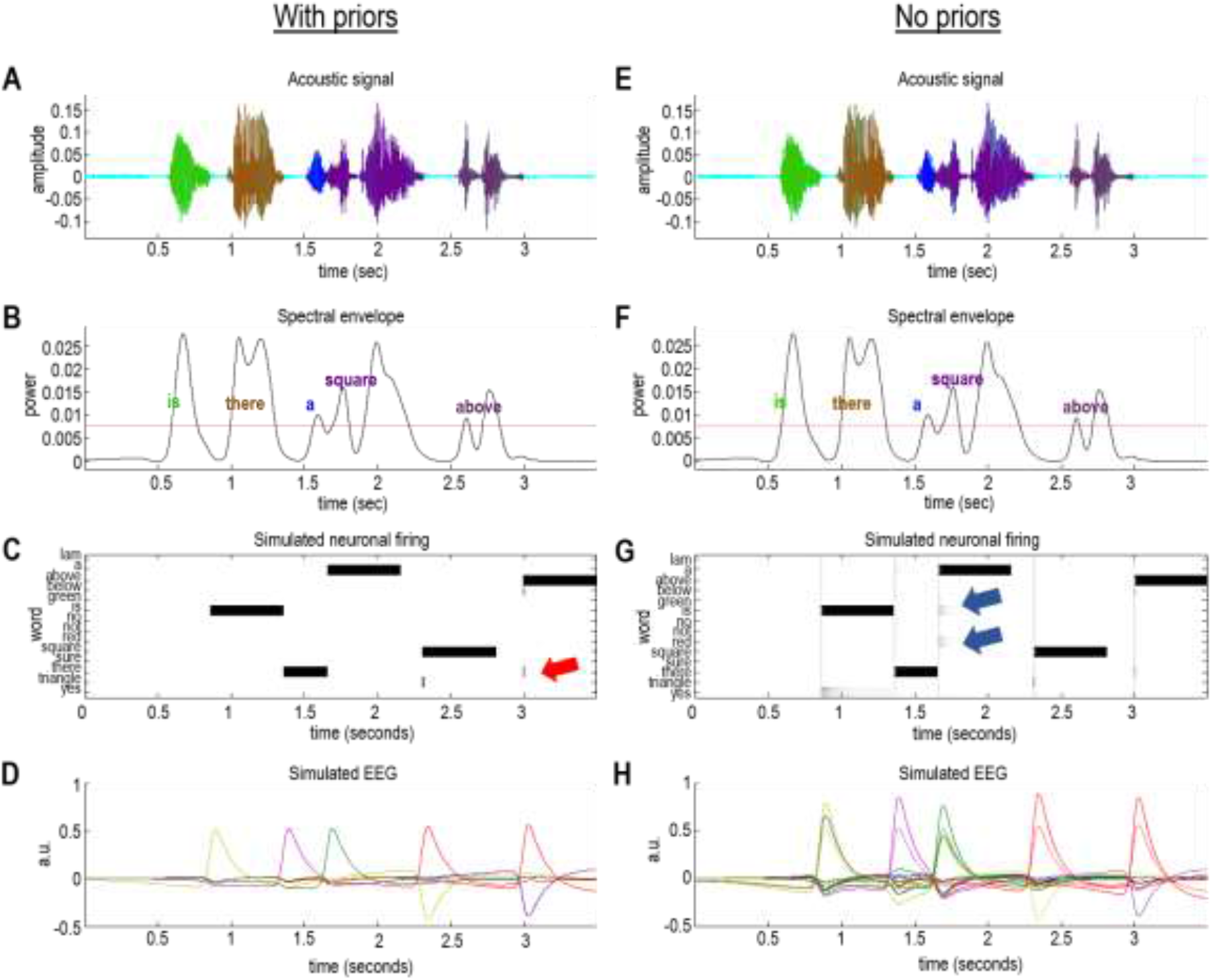
The role of priors in a word recognition: This figure uses the same format as Figure 6. In this example, the spoken sentence was “Is there a square above?” The left panel (A–D) shows the results of segmentation and word recognition under informative priors about the possible words. In other words, for each word in the sequence, a small number of plausible options were retained for inference. For example, the word “above” could have been “below” or “there”, as shown by the initial neuronal firing in Panel C at the end of the last word (red arrow). The right panel (E–H) shows exactly the same results but in the absence of any prior beliefs. The inference is unchanged; however, one can see in the neuronal firing (Panel G) that other candidates are competing to explain the acoustic signal (e.g., blue arrows). The key observation is that the resulting uncertainty—and competition among neuronal representations—is expressed in terms of an increased amplitude of simulated electrophysiological responses. This can be seen by comparing the simulated EEG trace in Panel H—in the absence of priors (solid lines)—with the equivalent EEG response under strong priors (solid lines in Panel D, reproduced as dashed lines in Panel H). In this example, there has been about a 50% increase in the amplitude of evoked responses. A more detailed analysis of the differences in simulated EEG responses is provided in Figure 8.

In the absence of precise prior constraints, the uncertainty associated with speech recognition is expressed as an increased amplitude of simulated electrophysiological responses. This can be seen most clearly by comparing the simulated electrophysiological responses in the lower right panel: the dotted lines reflect belief updating in the absence of specific priors, while the dashed lines are the same responses under informative priors. Figure 8 drills down on these differences by focusing on the responses to the third word. In so doing, the simulated waveform looks very much like a P300 that is frequently observed in electrophysiological studies (Donchin and Coles 1988, Morlet and Fischer 2014, Ylinen, Huuskonen et al. 2016). To understand this more formally, the next section explains how these simulated electrophysiological responses were derived and how they can be interpreted in terms of belief updating and Bayesian surprise.

**Figure 8.**
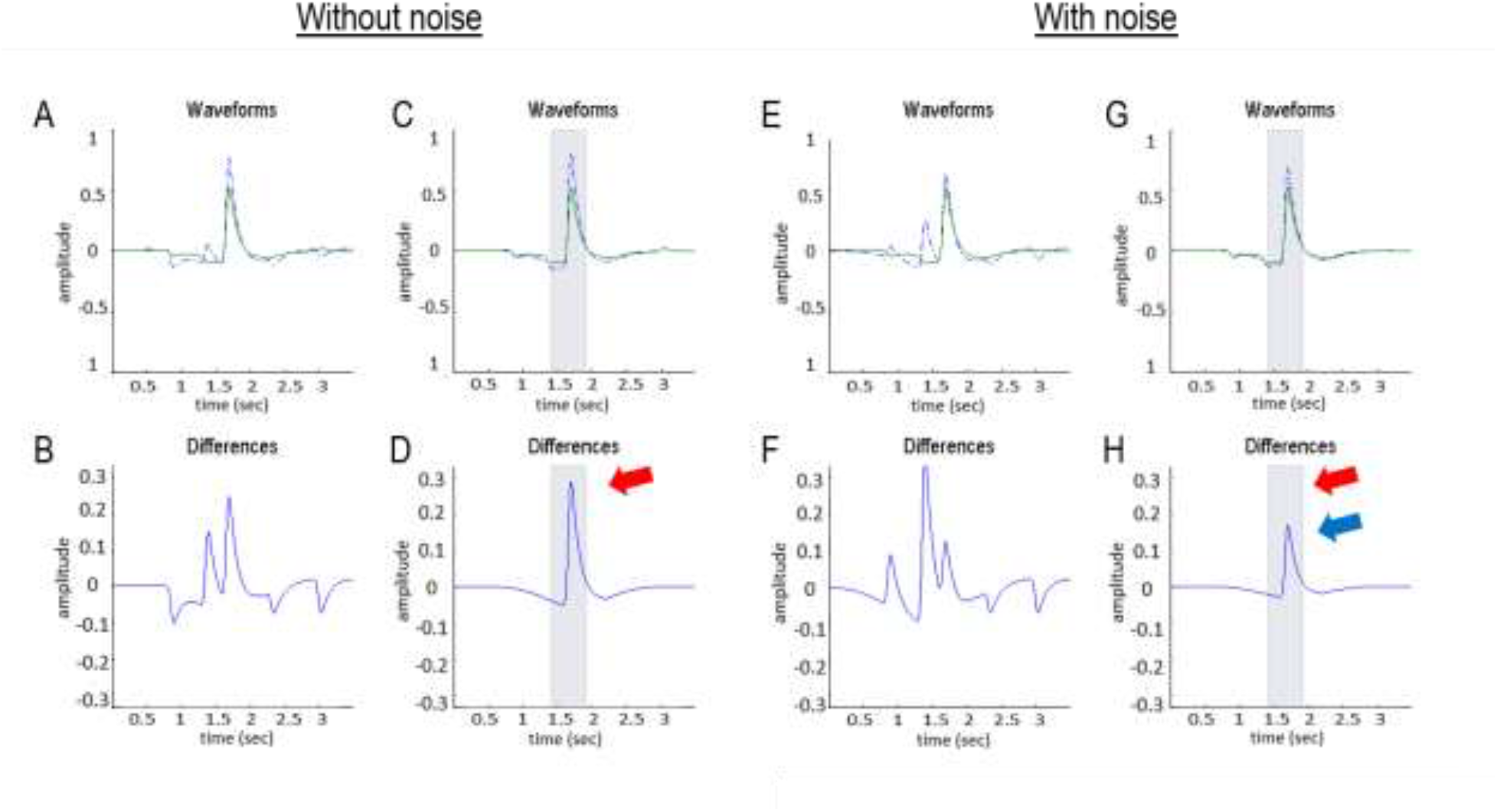
Mismatch responses and speech-in-noise: Panel A reproduces the results of Figure 7H, but focuses on the simulated electrophysiological responses of a single neuronal population responding to the third word (“a”). The upper row reports simulated responses evoked with (green lines) and without (blue dashed lines) priors (as in Figure 7), while the lower row shows the differences between these two responses. These differences can be construed in the spirit of a mismatch negativity or P300 waveform difference. Removing the priors over the third word (Panels C–D) isolates the evoked responses and their differences more clearly. The grey shaded area corresponds to a peristimulus time of 500 ms, starting 250 ms before the offset of the word in question. Assuming update time bins of around 16 ms means that we can associate this differential response with a P300. In other words, when the word is more surprising—in relation to prior beliefs about what will be heard—they evoke a more exuberant response some 300 ms after its offset. Panels E–H reports the same analysis with one simple manipulation; namely, the introduction of noise to simulate speech-in-noise. In this example, we doubled the amount of noise; thereby shrinking the coefficients by about a factor of half. This attenuates the violation (i.e., surprise) response by roughly a factor of two (compare difference waveform in Panel D without noise—red arrows—with the difference waveform in Panel H without noise—blue arrow). Interestingly, in this example, speech-in-noise accentuates the differences evoked in this simulated population when the word is not selected (i.e., on the previous word). The underlying role of surprise and prior beliefs in determining the amplitude of these responses is addressed in greater detail in the final figure.

To conclude this section, we will use this example to illustrate the fidelity of recursively generating and recognising words, under this generative model. Figure 9 shows the segmentation and word recognition following the presentation of the sentence above (“is there a square above”), without priors. The sentence was then generated using the recognised lexical, prosodic and speaker attributes. The synthetic speech was then presented to the active listening scheme, to recover the original utterance. This shows that the scheme can understand itself and perform rudimentary speech repetition. More formally, it illustrates the validity of the amortised inversion scheme.

**Figure 9.**
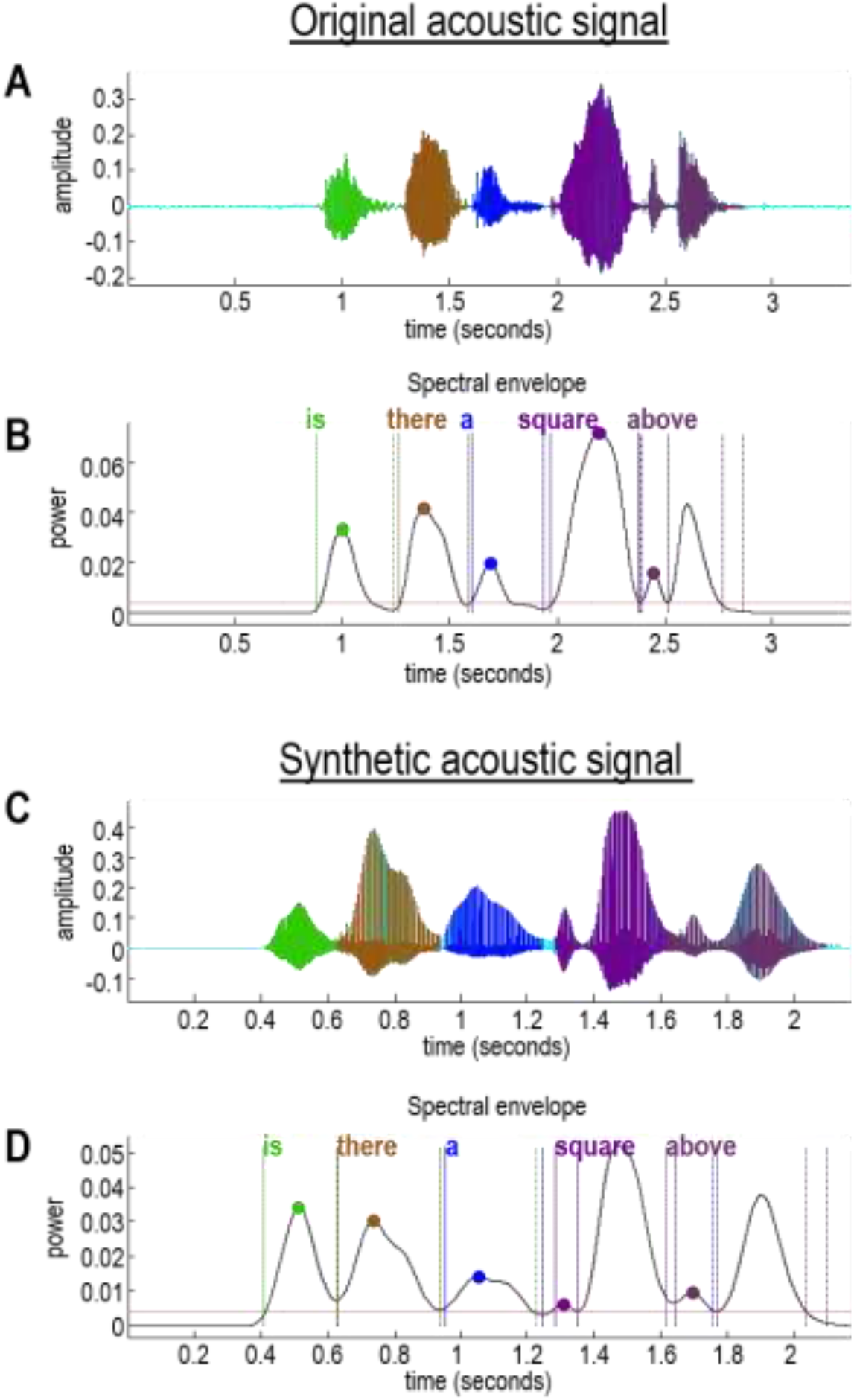
Recursive recognition and generation: The upper part of this figure shows the recognition of words (Panel B) contained within an acoustic signal (Panel A). Here, the acoustic signal is parsed into the words “is there a square above”. The corresponding lexical states can be used to synthesise a new acoustic signal (Panel C) containing the same words. Here, we inverted the model a second time, to recover the words contained within the synthetic acoustic signal (Panel D). Happily, the recovered words from the synthetic signal (Panel D) match those from the original signal (Panel B).

## Predictive validity: Belief updating and neurophysiology

Figure 8 suggests that belief updating during word recognition depends sensitively on prior beliefs and implicit differences in the confidence with which a particular word is inferred. Here, we pursue the predictive validity of this active listening formulation, by looking in greater detail at belief updating under the model. In doing so, we highlight qualitative similarities to canonical violation responses measured with EEG and MEG that are well-established in the empirical literature (as discussed in more detail below). In brief, the message of this section is that evoked or induced responses in the brain will increase in proportion to the degree of belief updating following sensory input.

Generally speaking, the idea that belief updating may underpin vigorous neuronal responses to surprising sensations is broadly consistent with experimental observations. Under predictive coding models of auditory perception, the mismatch negativity has been considered in light of precision weighted prediction error responses (Garrido, Kilner et al. 2009, Wacongne, Changeux et al. 2012, Heilbron and Chait 2018). In this literature, the mismatch negativity is related to deviants in elementary acoustic events, such as frequency (Näätänen, Gaillard et al. 1978, Giard, Lavikahen et al. 1995, Jacobsen, Schröger et al. 2003), intensity (Näätänen, Gaillard et al. 1978, Giard, Lavikahen et al. 1995, Jacobsen, Horenkamp et al. 2003), or timbre (Tervaniemi, Ilvonen et al. 1997, Tervaniemi, Winkler et al. 1997, Toiviainen, Tervaniemi et al. 1998)—and its amplitude covaries with the probability of a deviant (Picton, Alain et al. 2000, Sato, Yabe et al. 2000, Sato, Yabe et al. 2003). Mismatch negativity responses have also been recorded in the context of spoken phonemes (Dehaene-Lambertz 1997, Näätänen, Lehtokoski et al. 1997). In the current framework, precision weighted prediction errors induced by acoustic deviations reflect the surprise and concomitant belief updating induced by heard (spoken) words. At a slightly longer latency, reorientation responses could also be construed as a reflection of belief updating at higher levels of hierarchical inference. For example, the P300 has been proposed to reflect contextual violations (Donchin and Coles 1988) and the N400 has been proposed to reflect semantic violations (Kutas and Hillyard 1980, Kutas and Hillyard 1984, Van Petten, Coulson et al. 1999, Kutas and Federmeier 2000). The whole field of repetition suppression and adaptation in functional magnetic resonance imaging rests upon exactly the same notion; namely, an attenuation of neuronal responses that induce less belief updating, in virtue of being predictable or repetitious (Larsson and Smith 2012, Grotheer and Kovács 2014).

In the current simulations, our agenda is to identify generic principles that may underpin neuronal responses to surprising sensations under active listening. Our goal was not to simulate any particular type of ERP component, but merely to observe belief updating in the current framework. In the discussion section, we visit the finer details of the mismatch negativity and later endogenous (e.g., P300, N400) responses, which would be interesting avenues for future work. An advantage of the current setup is that we can expand upon the qualitative explanation for violation or surprise related responses using explicit, quantitative simulations.

If we take the average change in depolarisation under expected firing rates (after belief updating), we recover a quantity that scores the degree of belief updating (see Appendix 4 for details)—a quantity that emerges in many guises in different disciplines. For example, in statistics, it is known as the *complexity* (see equation A.18), which scores the departure from prior beliefs required to provide an accurate account of some data (Penny 2012). In the visual neurosciences, this quantity is known as *Bayesian surprise* (Schmidhuber 1991, Itti and Baldi 2009) that underwrites the *salience* or epistemic affordance of locations in the visual scene that attract saccadic eye movements (Parr and Friston 2017). In robotics, this quantity is known as *intrinsic motivation*; namely the *information gain* associated with a particular move or action (Ryan and Deci 1985, Oudeyer and Kaplan 2007). In short, we have a link between the information theoretic quantity that reflects the degree of Bayesian belief updating and the average neuronal responses that perform belief updating.

There are a number of reasons that one might consider this a sensible predictor of evoked responses in the brain, above and beyond the idealised dynamics described above. These reasons rest upon the statistical physics of belief updating in any sentient system making inferences about external states of affairs. The technical back story to active inference—that is, the free energy principle—allows one to associate the degree of belief updating and implicit changes in variational free energy in terms of a thermodynamic potential (Landauer 1961, Bennett 2003, Friston 2013). This means that for an ensemble of neurons (or neuronal processes) belief updating can be translated directly into thermodynamic free energy. The corresponding thermodynamic cost of belief updating may be reflected in nearly every sort of electrophysiological neuroimaging measurement. For example, the excursions of transmembrane potentials from their Nernst equilibrium in EEG (c.f., a mismatch negativity amplitude). Similarly, in fMRI, activations may reflect the metabolic costs of belief updating (Attwell and Iadecola 2002).

The second line of argument is based upon the common sense observation that, in the absence of an informative sensory cue, there can be no belief updating and no complexity cost or accompanying thermodynamic cost (Sengupta, Tozzi et al. 2016). In this instance, there will be, clearly, no evoked or induced response. This argument further suggests that the precision of continuous sensory (e.g., auditory) signals will determine the degree of belief updating and related violation responses, such as the mismatch negativity. In speech perception, reduced precision could correspond to speech-in-noise, for which this model predicts an attenuation of mismatch responses as noise levels increase. The basis of this effect rests upon the estimation of random fluctuations in sensory cues that, under predictive coding, shrink the posterior expectations of the lexical coefficients towards their prior mean.

If we revisit the results in Figure 6 and Figure 7, and compare responses evoked with and without priors, it is immediately obvious that, on average, evoked responses in the absence of (accurate) priors have a larger amplitude. This is sensible because priors that are congruent with the words presented mean that the belief updating has a smaller complexity cost because the prior is closer to the posterior. In other words, there is less information gain because the (synthetic) subject already had accurate prior beliefs about the lexical content of the spoken words.

To illustrate the sort of effect more quantitatively, we repeated the simulations reported in Figure 7 but introduced uncertainty about the third word by relaxing its priors. This allowed us to introduce differences in belief updating, from word to word, and show that simulated neuronal responses vary monotonically with information gain or Bayesian surprise. Figure 10 reports the results of this numerical analysis in terms of the variance of depolarisation over neurons encoding lexical expectations (blue line in the second panel) and the corresponding Kullback-Leibler divergence (red bars). Their monotonic relationship is apparent (see the third panel), although the relationship is not perfect due to filtering the simulated EEG data and our *ad hoc* measure of neuronal responses. At the (coarse-grained) level of the current treatment, this can be regarded as a simulation of neuronal responses to Bayesian surprise at a fairly high level in the auditory hierarchy (encoding the lexical content of a word).

**Figure 10.**
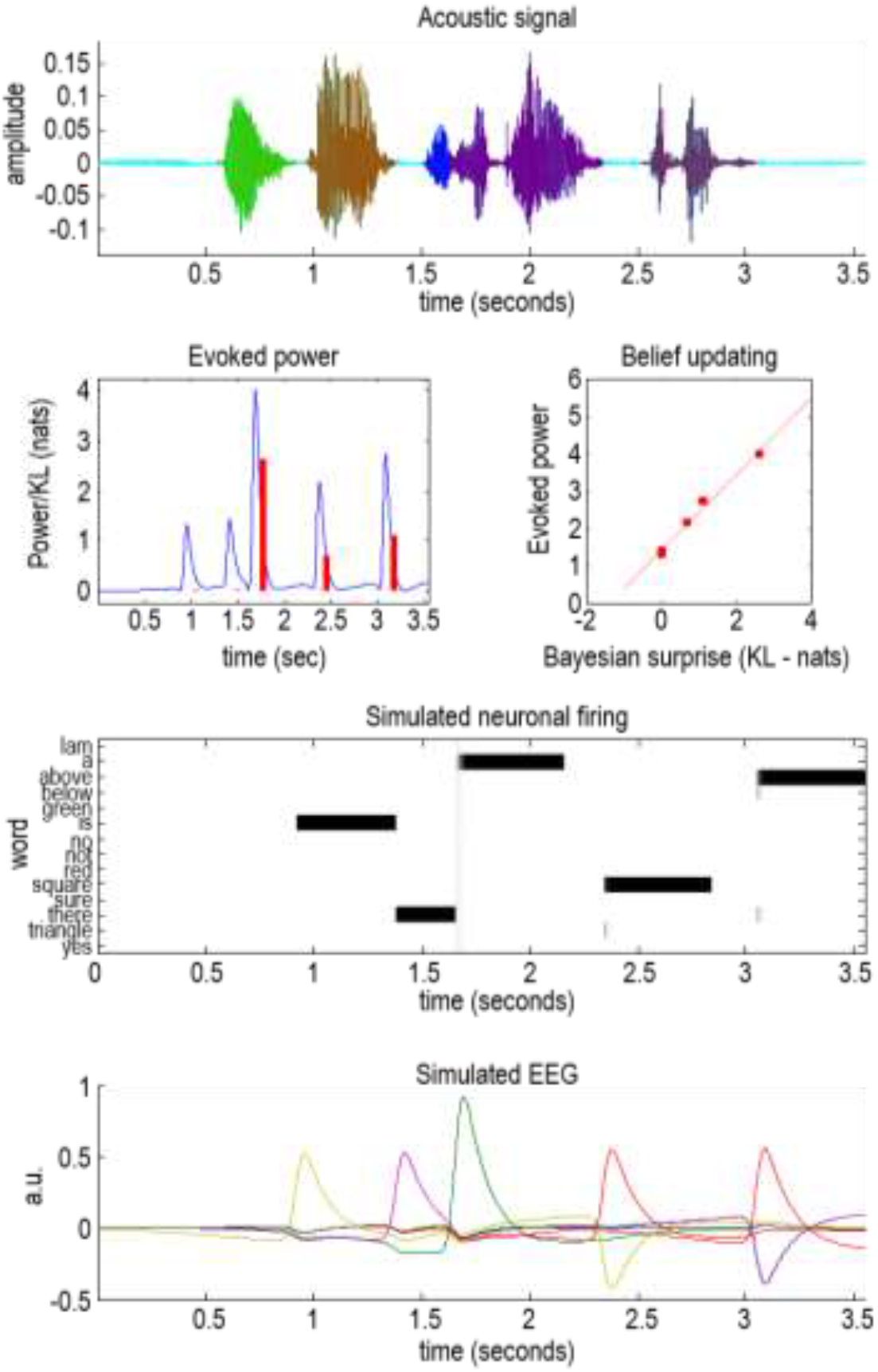
Bayesian surprise and evoked responses: this shows the same results as in Figure 7 but after removing priors from the third word (“a” in blue). The result is a more vigorous simulated event related response after the onset of the third word (green line in the bottom panel). A simple measure of these surprise-related responses can be obtained by taking the variance of the (simulated) responses over all populations as a function of time (c.f., evoked power). This is shown in the second panel as a solid blue line (normalised to a maximum of four arbitrary units). The red bars correspond to the degree of belief updating or Bayesian surprise, as measured by the KL divergence between prior and posterior beliefs after updating. The key conclusion from these numerical analyses is that there is a monotonic relationship between the evoked power and Bayesian surprise, as shown by the nearly linear relationship between Bayesian surprise and the maxima of evoked power in the third panel. In short, the greater the Bayesian surprise, the greater the belief updating and the larger the fluctuations in neuronal activity.

With this characterisation of mismatch responses, we can now return to the effect of noise, which highlights a key feature of active listening—that the quality of sensory evidence affects the magnitude of belief updating. In Figure 8, noise was simulated by decreasing the prior precision associated with the lexical coefficients at the auditory level of inference (namely, the prior precision in Equation A.20). This manipulation attenuates the mismatch or surprise response because the degree of belief updating has been reduced. The attenuation arises because there is less confidence placed in the evidence ascending from lower (sensory) levels of auditory processing. In other words, the attenuation of belief updating (and mismatch responses) in Figure 8 arises because the posteriors have been moved closer to the priors. This contrasts Figure 7, in which belief updating and mismatch responses were attenuated by one moving the priors closer to the posteriors. In subsequent work, we will revisit the effects of manipulating speech-in-noise—and prior beliefs—to demonstrate their effects empirically and, crucially, how they interact in the genesis of difference waveforms. For the purposes of this paper, the basic phenomenology illustrated above will be taken as a validation of the belief updating scheme by appealing to the literature on the canonical mismatch and violation responses of this sort.

## Discussion

Active listening considers the enactive synthesis or inference that might underwrite the recognition—and generation—of spoken sentences. The notion of *active listening* inherits from active inference, which considers perception and action under a universal imperative—to maximise the evidence for our (generative) models of the world. Here, the ‘active’ component is the (covert) parsing of words from a continuous auditory signal. Active listening entails the selection of internal actions (i.e., placement of word boundaries) that minimise variational free energy. Practically, word boundaries are selected so as to minimise surprise or maximise the evidence for an internal model of word generation. We have described the formal basis of this kind of active listening, using simulations of speech recognition to establish its face validity in behavioural terms. We then considered predictive validity, in terms of neuronal or physiological responses to violations and surprise, of the sort associated with the mismatch negativity, P300, and N400.

In treating the segmentation of a continuous sensory stream into meaningful words as an active sensing problem, we imagine that several segmentation operations are applied by the auditory system in parallel and the interval that maximises model evidence or marginal likelihood (i.e., minimises variational free energy) is selected for further hierarchical processing. From the perspective of hierarchical Bayesian inference, this follows the usual way of mapping from posterior density estimates, based upon continuous signals, to posterior beliefs about the discrete causes of those signals. This is generally cast in terms of Bayesian model selection. In other words, selecting some discrete explanation or hypothesis for the data that is most consistent with the estimated parameters of a generative model at the lower (sensory) level (Friston, Parr et al. 2017). The twist here is that this model selection has been framed in terms of action selection by treating the selection of word boundaries as an active process.

The generative model of word production that we considered has been stripped down to its bare essentials. More complex models could be conceived that synthesise more natural speech. Expanding the parameter space would not only allow it to produce more natural speech, but also allow the model to explain more domains of auditory production and perception. We discuss some of these possibilities in the discussion that follows. Nevertheless, we have demonstrated with this simplified generative model that inversion of the model—which corresponds to speech recognition—is associated with belief updating that makes plausible predictions for neuronal dynamics. In this paper, we produced quantitative simulations of electrophysiological responses and showed that they depend on the prior knowledge of the listener—a phenomenon that has commonly been observed in human speech perception (Marslen-Wilson 1975, Marslen-Wilson and Welsh 1978, Cole, Jakimik et al. 1980, Mattys and Melhorn 2007, Mattys, Melhorn et al. 2007, Kim, Stephens et al. 2012).

In borrowing ideas from active vision, we highlight parallels by which the brain could plausibly accumulate evidence among sensory modalities. The covert actions considered in this paper (i.e., the placement of word boundaries) follow in the spirit of overt (motor or autonomic) actions that have been used to simulate saccadic searches of the visual scene (Mirza, Adams et al. 2016, Parr and Friston 2017). We discuss the relationship between covert and overt actions in greater depth below. Intuitively, sensory observations in the auditory and visual modalities may appear to differ because speech unfolds over time, whereas visual experiments frequently use static stimuli that are spatially distributed. However, many parallels can be drawn between cortical processing in these modalities (O’Leary 1989), consistent with findings that sensory cortices can reorganise and subsequently process inputs from a different sensory modality (Sur, Garraghty et al. 1988, Shiell, Champoux et al. 2015). Shamma and colleagues (Shamma 2001, Shamma, Elhilali et al. 2011) propose a unified computational framework for auditory and visual perception, suggesting that the neural processes proposed for vision could also operate in auditory cortex. In short, this is based on the idea that the cochlea transforms temporally unfolding sound into spatiotemporal response patterns early in auditory processing. In other words, this is a ‘spatial’ view of auditory processing. Under this view, the computations for analysing auditory signals in time could be similar to the computations used for analysing visual signals in space; e.g., (Bar, Kassam et al. 2006).

### Active listening and Bayesian surprise

Selecting intervals containing auditory cues that minimise free energy (i.e., maximise marginal likelihood or model evidence) follows from the basic premise of the free energy principle; namely, both action and perception are in the game of self-evidencing (Hohwy 2016). Having said this, there is something unique about the particular selective process (which are implicit in Equation A.19) that distinguishes it from overt actions, such as moving one’s head or making visual saccades to a location in a visual scene. This is because the corresponding selection of ‘where to look next’ is based upon anticipated data that would be sampled if one looked ‘over there’. However, predictive coding (in some amortised form) of speech segmentation here is based on evidence *that has already accumulated* under different interval or segmentation schemes. In other words, there is a distinction between overt actions—such as moving one’s eyes or moving one’s head—which changes observations in the future, and covert actions—such as covert visual attention, or selecting a particular segmentation of speech—which is based on sampling current observations. In the case of these covert actions, the sensory evidence (and subsequent posterior) can be computed explicitly to evaluate the free energy expected under a particular interval choice. In contrast, expected free energy based on overt actions has to be averaged under predicted sensory outcomes—known technically as a posterior predictive density. This means that evaluating the *free energy* for particular speech segmentation intervals is much simpler than evaluating the *expected free energy* under a posterior predictive density, conditioned upon a particular overt action. It is useful to bear this distinction in mind because it can resolve some apparent paradoxes.

These paradoxes pertain largely to the question: does active inference minimise or maximise Bayesian surprise? In the current setting, covert actions associated with speech segmentation minimise Bayesian surprise, because Bayesian surprise relates to the complexity (i.e., cost) associated with belief updating based on current observations. In other words, because the free energy associated with covert actions can be evaluated explicitly, a listener can choose the covert action that requires the least belief updating (i.e., that is closest to their priors), but still provides an accurate explanation for the auditory observations. This leads to a conceptualisation in which neuronal dynamics and implicit message passing aim to explain sensory input with minimal complexity and, therefore, minimum accompanying thermodynamic cost (Sengupta, Stemmler et al. 2013). On this view, large mismatch or violation responses indicate that an accurate explanation for sensory inputs required a costly update to posterior beliefs.

The situation flips for overt actions, for which action selection depends on *expected* free energy—which is evaluated on the basis of predicted (i.e., unknown) outcomes in the future. Future sensory outcomes are random (i.e., unknown or hidden) variables and active inference maximises expected Bayesian surprise, which corresponds to expected information gain. In other words, it reflects the reduction in uncertainty in how the world is sampled. Actions that maximise Bayesian surprise will lead to the greatest reduction in uncertainty. This is why *expected* Bayesian surprise has to be maximised when selecting actions, where it plays the role of epistemic affordance (Parr and Friston 2017). As noted above, this is an important imperative that underwrites uncertainty reducing, exploratory behaviour; known as intrinsic motivation in neurorobotics (Schmidhuber 2006) or salience when ‘planning to be surprised’ (Sun, Gomez et al. 2011, Barto, Mirolli et al. 2013). An intuitive way of thinking about whether surprise should be maximised or minimised is to appeal to the analogy of scientific experiment. We may attempt to analyse empirical data that we have collected in a way that minimises how surprising it appears; for example, by giving greater weight to hypotheses consistent with our measurements. Having done so, we may want to design a future experiment, which would aim is to collect data that will tell us something new; in this case, we should design an experiment that we expect to maximise our (Bayesian) surprise (a.k.a., information gain).

In future work, we will expand upon this distinction by using the current model to simulate conversations. The act of speaking is an overt action, and the basic principle of conversational turn taking has been simulated using active inference in the setting of bird song (Friston and Frith 2015). We hope to combine the current active listening implementation with an agent who is able to ask questions. In brief, the agent will actively listen to speech by *minimising* Bayesian surprise at the level of word recognition considered in this paper, and select words to speak (i.e., overt actions, here in the form of questions) that *maximise* expected Bayesian surprise to maximise information gain (i.e., resolve uncertainty). This leads to a first principle account of language ‘understanding’ that can be described in terms of self-evidencing: namely, minimising free energy through belief updating, and planning to take actions that minimise expected free energy.

Although evaluating the free energy of alternative data features (i.e., segments) that have already been sampled is more straightforward than evaluating the expected free energy when planning how to sample data, it is not as straightforward as reflexive action; e.g., (Adams, Shipp et al. 2013). Reflexive or elementary action, under active inference, changes the sensory data solicited, e.g., the stretch receptor signals that are attenuated by classical motor reflexes. However, this kind of reflexive action does not change internal brain states or the posterior beliefs that they parameterise. This means that the only part of free energy that can be minimised directly is the accuracy term (Equation A.18). This is why it is sufficient to minimise interoceptive and proprioceptive prediction errors when accounting for autonomic and motor action; very much along the lines of the equilibrium point hypothesis (Feldman and Levin 1995) and the passive movement paradigm (Mohan and Morasso 2011). However, in the active listening framework proposed here, the situation is a little more involved. This is because hierarchical inference means that committing to one data feature (i.e., interval) or another will change posterior beliefs. This means that to comply with the free energy principle, it is necessary to select data features (i.e., intervals) that not only maximise accuracy but also minimise complexity. This entails a more nuanced form of action selection, in virtue of the fact that it requires the (covert) selection of data features that have been (overtly) acquired. Even though the data have already been acquired, and selecting different data features does not change the auditory outcomes (acoustic timeseries), these processes are nevertheless ‘active’ from our perspective, because the agent has an epistemic imperative to sample auditory outcomes in a way that reduces uncertainty. In other words, the agent is in charge of the *data features* (i.e., segmentation). Thus, we can think of speech segmentation as a kind of action that is internal or attentional, related to how the acoustic timeseries is covertly sampled. The framework we have introduced in this paper highlights that— mathematically—these covert actions can be considered in a similar way as overt actions.

### Acoustic envelope and spectral fluctuations

Under active listening, the implicit generative model of an envelope, which is used to create a repertoire of intervals from which to select, is distinct from the spectral fluctuations (i.e., formant frequencies) generated by latent states (i.e., lexical and prosody). This formulation of speech recognition may explain why there are ‘envelope following responses’ in distinct parts of the auditory system, whose functional architecture can be distinguished from the tonotopic mapping of auditory cortex per se (Easwar, Purcell et al. 2015, Braiman, Fridman et al. 2018). This leads to an interesting picture of how the brain thinks words are generated that echoes the distinction between ‘what’ and ‘where’ in the visual hierarchy (Ungerleider and Haxby 1994). In other words, there may be a homologous distinction between ‘what’ and ‘when’ in the auditory system that manifests as an anatomical separation of the pathways inferring ‘what’ is being spoken (i.e., tonotopic predictions and representations) and when this content is deployed (i.e., envelope following responses) (Romanski, Tian et al. 1999, Alain, Arnott et al. 2001). From the point of view of word generation, these two streams converge to generate the correct formants at the correct time. From the point of view of recognition or generative model inversion; this would imply a functional segregation of the sort seen in other modalities (Ungerleider and Haxby 1994, Friston and Buzsaki 2016); for example, the segregation into dorsal and ventral streams – or, indeed, parvocellular and magnocellular streams (Zeki and Shipp 1988, Nealey and Maunsell 1994). Interestingly, this sort of segregation into ‘what’ and ‘how’ pathways has already been proposed for the auditory system (Kaas and Hackett 1999, Belin and Zatorre 2000).

### Active listening and electrophysiological responses

In a general sense, we have shown that belief updating under active listening qualitatively resembles physiological responses to violations and surprise that are already in the literature. Our goal was not to simulate any particular type of ERP component or the empirical results from any particular study, but rather to explore belief updating in an artificial agent whose goal is to generate and/or recognise speech. So, can we interpret this belief updating in light of particular ERP responses?

One canonical violation response is the mismatch negativity. The mismatch negativity is observed in classic ‘oddball’ paradigms (Garrido, Kilner et al. 2009), in which a deviant sound follows a sequence of sounds that all share a particular acoustic property. Mismatch negativity responses have been observed when a sound deviates in frequency (Näätänen, Gaillard et al. 1978, Giard, Lavikahen et al. 1995, Jacobsen, Schröger et al. 2003), intensity (Näätänen, Gaillard et al. 1978, Giard, Lavikahen et al. 1995, Jacobsen, Horenkamp et al. 2003), or timbre (Tervaniemi, Ilvonen et al. 1997, Tervaniemi, Winkler et al. 1997, Toiviainen, Tervaniemi et al. 1998) from preceding stimuli. Crucially, the mismatch negativity has recently been interpreted in terms of predictive coding—specifically, it has been assumed to reflect precision weighted prediction errors (Garrido, Kilner et al. 2009, Wacongne, Changeux et al. 2012, Heilbron and Chait 2018)—which relates nicely to the current framework. The finding that the amplitude of the mismatch negativity covaries with the probability of a deviant (Picton, Alain et al. 2000, Sato, Yabe et al. 2000, Sato, Yabe et al. 2003) is consistent with the idea that it reflects belief updating. Most previous studies of the mismatch negativity have used basic auditory stimuli, such as artificial pure or complex tones; it is therefore assumed to reflect deviations to low-level acoustic properties, rather than processes that are specific to speech. Nevertheless, observations of the mismatch negativity during phoneme perception (Dehaene-Lambertz 1997, Näätänen, Lehtokoski et al. 1997) can be interpreted as reflecting acoustic violations that occur within speech.

The P300 is often observed in similar ‘oddball’ settings as the mismatch negativity (Polich 2007). It has a longer latency than the mismatch negativity and has been related to higher-level context violations (Donchin and Coles 1988). It could, therefore, be interpreted as reflecting belief updating when the listener’s context changes. In the domain of speech, the P300 has been associated with word frequency (Polich and Donchin 1988).

The N400 is commonly observed in response to meaningful speech, and has also been associated with word frequency (Kutas and Hillyard 1984, Van Petten and Kutas 1990, Van Petten, Coulson et al. 1999). Kutas and Hillyard (Kutas and Hillyard 1984) found that the amplitude of the N400 was inversely correlated with a word’s cloze probability—that is, participants’ ratings of the probability that a particular word would come at the end of the sentence in question. They found that the same effect transferred to words that were semantically related to high-probability words. They, therefore, concluded that the N400 relates to semantic activation. Modulations of N400 responses have been reported in a variety of semantic contexts (reviewed by (Kutas and Federmeier 2000))—including sentence-final words, the semantic congruency of words that occur mid-sentence, and the semantic relatedness of word pairs—and has been shown to build up as the semantic context becomes increasingly constrained throughout a sentence. Syntactic violations do not elicit an N400 response (Kutas and Federmeier 2009), but instead evoke a P600 (Osterhout and Holcomb 1992, Friederici, Hahne et al. 1996, Kuperberg, Sitnikova et al. 2003).

An N400-like negativity, termed the frontocentral negativity (‘FN400’) has been related to speech segmentation by transitional probabilities (Balaguer, Toro et al. 2007, Cunillera, Càmara et al. 2009, François, Cunillera et al. 2017). For example, stronger FN400 responses were elicited from acoustic signals that comprised strong statistical relationships between syllables than syllables that were selected randomly (François, Cunillera et al. 2017). The FN400 also appears to increase in amplitude as the segmentation process becomes more prominent as new words are learned (Balaguer, Toro et al. 2007, Cunillera, Càmara et al. 2009).

Speech segmentation by prosodic cues has been associated with a different ERP: the closure positive shift (CPS) (Steinhauer, Alter et al. 1999). The closure positive shift is evoked around the time of a prosodic boundary, and has been reported to last until the onset of the next word (Bögels, Schriefers et al. 2011). It has been found in several different languages (see (Bögels, Schriefers et al. 2011) for a review) and even in hummed speech (Pannekamp, Toepel et al. 2005), which has no lexical content.

So, which level of processing does belief updating in the current scheme reflect? This level could be intermediate between lower acoustic levels at which a mismatch negativity is generated, and the kind of violation responses associated with a change in context or semantics. Possibly, this could be something like the phonological mismatch negativity, which has been interpreted as reflecting acoustic-phonetic processing in response to the initial phoneme of a spoken word, occurring 270–300 ms after onset (Connolly, Phillips et al. 1992). Connolly and Phillips (Connolly and Phillips 1994) observed the phonological mismatch negativity when the final word of a sentence was semantically congruent, but the word (and the initial phoneme) differed from the word with the highest Cloze probability. An N400 was not observed in this condition and was instead observed when the word was semantically incongruent. Interestingly, the phonological mismatch negativity was not observed when a word was semantically incongruent, but the initial phoneme matched the word with the highest Cloze probability. These observations are consistent with the idea that the phonological mismatch negativity reflects acoustic-phonetic processing.

One advantage of the current framework is that it generates quantitative predictions that can be explicitly tested in future electrophysiological studies. The predictive validity we have considered here is a first step: the next step is to scrutinise the particular parameters of the simulation using empirical data. To study this in more detail, specific sequences of words and/or acoustic features could be posed to the model that generate particular violations. Belief updating in active listening—and, for comparison, parameters of other models (Aitchison and Lengyel 2017)—could be quantitatively compared to empirical electrophysiological results. This speaks again to future directions, in which the current framework will be extended to a hierarchical model that can simulate conversations. Speech has a deep temporal structure, with phrases evolving over longer time intervals than words or phonemes—and a more complete generative model of speech will have to incorporate this temporal hierarchy (Friston, Rosch et al. 2017). The idea of an interlocutor asking questions to resolve uncertainty relates to a higher-level semantic processing of speech—and violations of semantic expectations might be associated with later electrophysiological responses, such as the N400. Consistent with the types of hierarchies that have often been suggested based on empirical data (Kumar, Stephan et al. 2007, Ding, Melloni et al. 2015), a deep generative model implies that belief updating occurs at multiple time scales, and we anticipate that this will give rise to more structured ERPs that include contributions from later components.

### Background noise during active listening

In this paper, we simulated a simple case of speech-in-noise, in which we imposed random fluctuations (of constant amplitude) on the speech signal. We showed that noisier signals attenuate belief updating. We plan to extend this model to incorporate other types of noise, including fluctuating-amplitude maskers such as multi-speaker environments. This should allow one to investigate which aspects of the signal are most informative for minimising Bayesian surprise, when some parts of the signal (but not others) undergo energetic masking (Brungart 2001, Brungart, Simpson et al. 2001, Durlach 2006) or when informational masking (Durlach, Mason et al. 2003, Durlach, Mason et al. 2003, Kidd, R. Mason et al. 2007) comes into play. In other words, in the presence of noise, a listener needs to reduce their uncertainty about the words that were spoken by deciding which attributes of the acoustic signal they should attend to.

One problem that the current segmentation algorithm would face—when adding background noise to speech—is that envelope minima may not always be present at word boundaries. In human listeners, segmentation at envelope minima could be achieved based on envelope following responses. Indeed, the magnitude of envelope following responses (i) has been linked to speech intelligibility in humans (Drullman 1995, Muralimanohar, Kates et al. 2017, Vanthornhout, Decruy et al. 2018), (ii) is greater for attended than unattended speakers (Ding and Simon 2012, O’Sullivan, Power et al. 2014), and (iii) can be reconstructed from measurements of brain activity (Pasley, David et al. 2012, O’Sullivan, Power et al. 2014). These envelope responses could, therefore, reflect the success of speech segmentation. Other cues to segmentation have been reported in the literature—and may be particularly important when background noise is present. These cues include durations: a lengthening of syllables at the end of words (Klatt 1975, Beckman and Edwards 1990), and possibly also the beginning (Lehiste 1960, Lehiste 1972, Oller 1973, Klatt 1976, Nakatani and Dukes 1977, Gow Jr and Gordon 1995). They also include a shortening of the middle portion of words (Lehiste 1973, Oller 1973, Harris and Umeda 1974, Klatt 1976). Other work has also reported metrical (stress) cues (Cutler and Norris 1988), allophonic variation (Christie Jr 1974, Nakatani and Dukes 1977, Gow Jr and Gordon 1995), and fundamental frequency contour (Ladd and Schepman 2003) as segmentation cues. Although the current algorithm of finding envelope minima was sufficient for the current simulations, these other cues could be implemented into active listening in other contexts in which segmentation may be particularly challenging. While the current implementation retrospectively places word boundaries, future work could also consider that word boundaries are somewhat predictable from the lexical statistics of the preceding sequences (Marslen-Wilson 1984)—for example, the offset of “trombone” may be predicted upon hearing “trom”, given it is the only valid ending to the word in English.

### Active listening and language production and perception

The active listening scheme can also be used as a foundation to gain a neuronal-level understanding of language production and perception behaviours. For example, engaging in a two-way dialogue (Kuhlen, Bogler et al. 2017), verbal fluency (Paulesu, Goldacre et al. 1997) and reading (Fiez and Petersen 1998, Landi, Frost et al. 2013, Taylor, Rastle et al. 2013); see (Price 2012) for a detailed overview. Previous investigations of these behaviours have been motivated by the desire to better understand the underlying neuropsychology (Aring 1963, Hodges, Patterson et al. 1992, Warburton, Price et al. 1999, Thiel, Habedank et al. 2005, Nardo, Holland et al. 2017, Hope, Leff et al. 2018). In other words, what are the causal mechanisms associated with (language) behavioural modifications following neurological disorders? Despite valiant efforts, none of the current computational accounts of language can fully explain these behaviours (Rueschemeyer, Gaskell et al. 2018): examples include Directions Into Velocities of Articulators model (Tourville and Guenther 2011), State Feedback Control model (Houde and Nagarajan 2011), and Hierarchical State Feedback Control model (Hickok 2014). Crucially, these approaches do not simultaneously account for higher-order language processing (semantic, syntactic, *etc*.) and lower level articulatory control (prosody, *etc*.); however, human language processing requires both. The active listening scheme presented here departs from previous approaches: it explicitly considers the segmentation of continuous signals (which come into play through the accuracy term in Equation (A.18) and relate to lower-level processing) and beliefs about the lexical content of those signals (key to the complexity term in Equation (A.18) and relating to higher-level language processing). Not only do these two aspects exist in the model, but they go hand-in-hand during word recognition. This makes the generative model described here a prime candidate for developing a mechanistic and neurobiologically plausible account of (healthy and impaired) language behaviour.

The idea that a generative model for speech generation can be inverted for the purpose of recognising speech touches upon a longstanding debate in the literature—are similar neural processes used to recognise speech, as those that are used to produce speech? This is an interesting question, and one that the current formulation does not address. Of relevance, the properties of spoken sentences that active listening uses to produce and recognise speech are acoustic (e.g., fundamental and formant frequencies) rather than biological (e.g., vocal chords and vocal tract) attributes (Guenther and Vladusich 2012). Thus, it does not necessarily follow from this framework that an individual who is unable to speak is unable to comprehend speech. On the contrary, we expect that an individual who is unable to speak could still generate an internal model that specifies the causes of spoken words, which they have learnt by perceiving speech. Whether the experience of producing speech contributes to the same model is an interesting question. In short, there may be an opportunity to examine how computational lesions to the model impair speech perception and production.

### Active listening and voice recognition

One strength of the current scheme is that it deals with both speech generation and recognition, and can be iteratively applied to recognise the lexical content of simulated speech (see Figure 9). The simulated speech that the model produces is discernibly artificial, but the key message here is that the model reduces the problems of speech generation and recognition to their necessary parameters. The generative model introduced in this paper lays the groundwork for a complete model of voice recognition. In other words, a model that infers *who* is speaking. The current model includes states for the speaker attributes of their average fundamental frequency and formant spacing. From a speech production perspective, a speaker’s fundamental frequency relates to the rate of vocal fold vibration (known as glottal pulse rate), and formant spacing is affected by the length and shape of the vocal tract—which are relatively fixed for a speaker, although can be modified slightly by changing the positions of the articulators, such as the tongue and lips. Previous research demonstrates that listeners use both fundamental frequency and speech formants to judge the identity of people who are familiar (LaRiviere 1975, Abberton and Fourcin 1978, Van Dommelen 1987, Van Dommelen 1990, Lavner, Gath et al. 2000, Lavner, Rosenhouse et al. 2001, Holmes, Domingo et al. 2018) and unfamiliar (Matsumoto, Hiki et al. 1973, Walden, Montgomery et al. 1978, Murry and Singh 1980, Baumann and Belin 2009, Gaudrain, Li et al. 2009). To extend the current model to recognise voices, the next step is to specify how combinations of fundamental and formant frequencies are used to infer speaker identity. From the perspective of the generative model, fundamental and formant frequencies are generated from hidden states that correspond to particular speakers. This approach differs from that proposed by Kleinschmidt and Jaeger (Kleinschmidt and Jaeger 2015), who assume that listeners construct a separate generative model for each talker they encounter. In the current implementation, we have focused on fundamental and formant frequencies, because these attributes are most prevalent in the voice recognition literature. However, they are not the only relevant speaker attributes (Cai, Gilbert et al. 2017, Holmes, Domingo et al. 2018). More complex models of voice recognition could incorporate additional speaker parameters, for example, relating to speaker-specific accent, stress, and intonation.

### Active listening and music

Finally, the generative and inversion schemes presented here could also form the basis for models of other complex auditory signals. Music, for example, shares several features with language (Patel 2010) and relies on partly overlapping brain networks (Musso, Weiller et al. 2015), which makes it a natural choice for future work. It is not difficult to imagine how the generative model in Figure 1 could be adapted to simulate music in an active listening framework. For example, somewhat akin to determining the correct onsets and offsets of word boundaries, we need to decide where a musical phrase—or longer section of music—begins and ends.

Recent empirical findings have shown that mismatch responses to unexpected musical sounds are larger in contexts with low than high uncertainty (Quiroga-Martinez, Hansen et al. 2019). This fits comfortably with the proposed explanation of evoked responses as reflecting Bayesian surprise or salience, which would be reduced when sensory signals are unreliable or imprecise. Since music is rich and multifaceted and relies greatly on statistical learning (Pearce 2018), it would be an ideal means to understand how neuronal dynamics change with uncertainty.

### Summary

In summary, this paper introduces active listening—a unified framework for generating and recognising speech. The generative model specifies how discrete *lexical*, *prosodic*, and *speaker* attributes give rise to a continuous acoustic timeseries. As the name implies, the framework also includes an active component, in which plausible segmentations of the acoustic timeseries—corresponding to the placement of word boundaries—are considered, and segmentation that minimises Bayesian surprise is selected. In the simulations presented here, we demonstrate that speech can be iteratively recognised and generated under this model. We show that the words that the model recognises depend on prior expectations about the content of the words, as is the case for human listeners, and that simulated neuronal responses resemble human electrophysiological responses. This work establishes a foundation for future work that will simulate human conversations, voice recognition, speech-in-noise, and music—and which we anticipate will provide key insights into neuropsychological impairments to language processing.

## Software note

The routines described in this paper are available as Matlab code in the SPM academic software: http://www.fil.ion.ucl.ac.uk/spm/. The simulations reported in the figures can be reproduced (and customised) via a graphical user interface by typing (in the Matlab command window) **DEM** and selecting appropriate (speech recognition) demonstration routines. The accompanying Matlab scripts are called **spm_voice_*.m**.

## Acknowledgements

The Wellcome Trust funded K.J.F. (Ref: 088130/Z/09/Z), E.H. (Ref: WT091681MA), and the Wellcome Centre for Human Neuroimaging (Ref: 203147/Z/16/Z), where this work was conducted. N.S. is funded by the Medical Research Council (Ref: MR/S502522/1). D.R.Q. is funded by the Danish National Research Foundation (Project number: DNRF117). T.P. is supported by the Rosetrees Trust (Award number: 173346).

## Appendices

## Appendix 1: The generative model

This appendix covers technical details of the generative model introduced in Figure 1. Figure 11 is designed to supplement Figure 1, and includes the equations corresponding to word generation (left column) and word recognition (right column). This section first provides a summary of the technical details of the generative model, then goes on to unpack each of the equations of the generative model in Figure 11. Although these may seem complicated for a non-technical reader, they are simply a sequence of non-linear transforms that specify the mapping from lexical, speaker, and prosody parameters to an acoustic timeseries.

**Figure 11.**
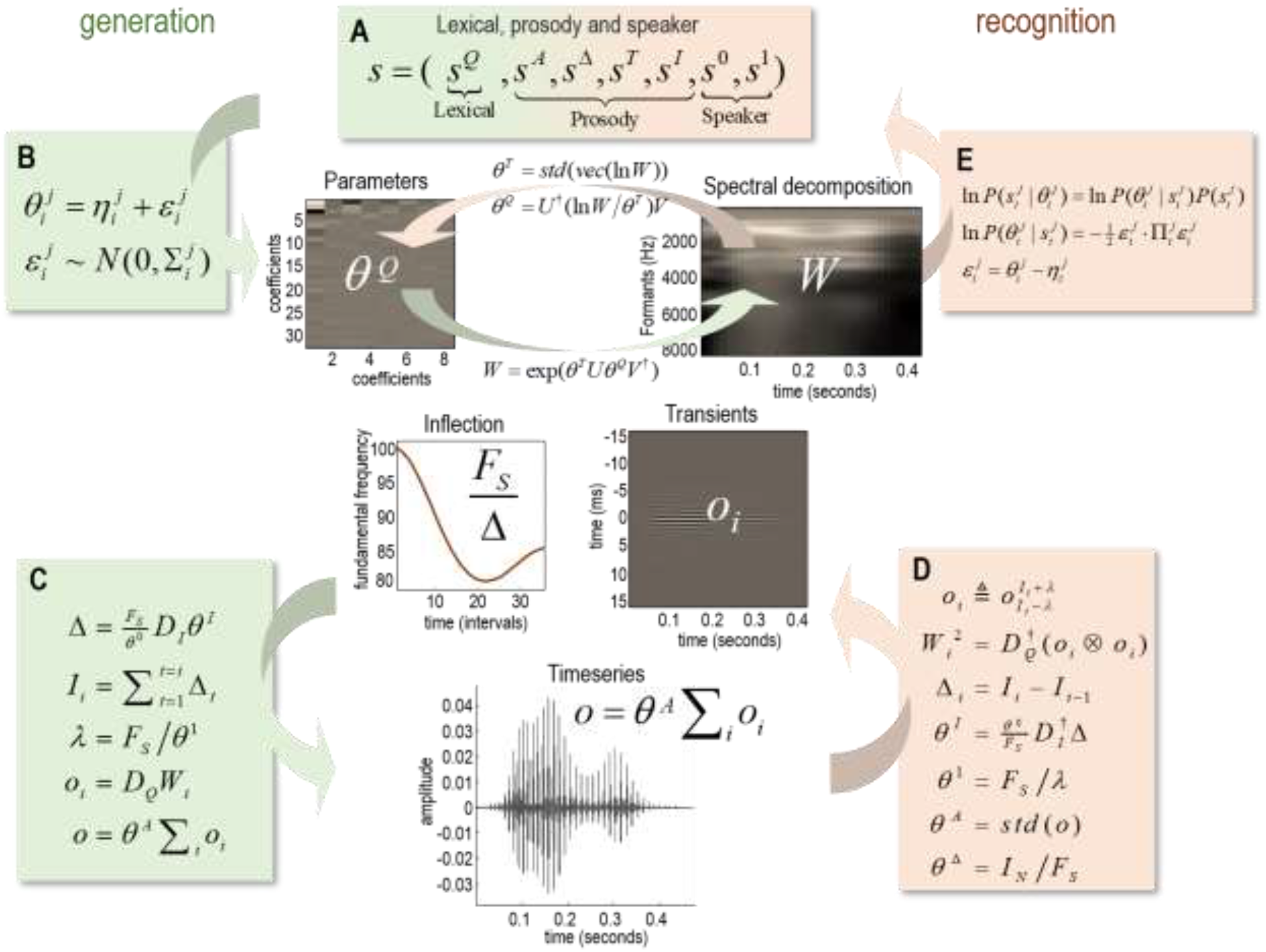
A generative model of a word. This figure illustrates the generative model from the perspective of word generation (green panels) and accompanying inversion (orange panels), which corresponds to word recognition. This model maps from hidden states (*s*; shown in box A), which denote the attributes of a spoken word (in this case lexical content, prosody, and speaker identity), to outcomes (*o*; shown in box C), which corresponds to the continuous acoustic timeseries. Box B shows how parameters are sampled for word generation. The centre panels illustrate the non-linear mappings between model parameters and the acoustic spectrum (i.e., time-frequency representation). Box C specifies how the transients are then aggregated to form a timeseries. Recognition (boxes D–E) corresponds to the inversion of the generative model: a given time series is transformed to parameterise the time-frequency representation (box D) by simply inverting or ‘undoing’ the generative operations. These parameters are used to evaluate the likelihood of lexical, prosody and speaker states (box E). The equations displayed in this figure are unpacked in the text.

In brief, each word (i.e., lexical item) is associated with a matrix of a discrete cosine transform coefficients (*θ^Q^*) that generate a time-frequency representation (*W*) of the spoken word (i.e., the spectrogram), when combined with speaker and prosody information. In this scheme, the lexical form and structure comprise a discrete cosine transform with 8 basis functions over time and 32 over formant frequencies (see Figure 11C). The number of basis functions was selected as a compromise between the quality of the generated acoustic timeseries and computational efficiency. Each column of the time-frequency representation generates a transient: thus, the number of transients corresponds to the number of columns in the time-frequency representation.

The transients are emitted at an instantaneous fundamental frequency, which is inversely proportional to the time intervals between successive transients (Δ_*i*_). These time intervals are stored in a fundamental interval variable (*I*). The instantaneous fundamental frequency is affected by the average fundamental frequency of the speaker (*θ*^0^), corresponding to their average *glottal pulse rate*. It also depends on a discrete cosine transform (*D*) based upon (three) coefficients (*θ^I^*) that encode inflection around the speaker’s average fundamental frequency (*θ*^0^): (1) the average fundamental frequency relative to the speaker average, (2) increases or decreases in fundamental frequency over time, and (3) the acceleration or deceleration of changes in fundamental frequency. The ensuing time-frequency representation is then multiplied by an inverse temperature (*θ^T^*) parameter, which affects the quality of the sound and can be thought of as a timbre parameter. Its exponential is, effectively, Fourier transformed to create a succession of transients that are deployed over fundamental intervals. The resulting timeseries is then scaled by an amplitude parameter (*θ^A^*) to furnish the final (continuous) acoustic timeseries.

In what follows, we unpack each of the equations in Figure 11, from the perspective of word generation (left column of Figure 11). Note that word generation simply involves a sequence of non-linear transformations, which specify the relationship between parameters and the acoustic timeseries.

Each discrete state generates a parameter that is sampled from a Gaussian distribution (Figure 11B) with a mean *η* and covariance *Σ*. The subscript notation indicates hidden state *j* and its *i*-th possible value:

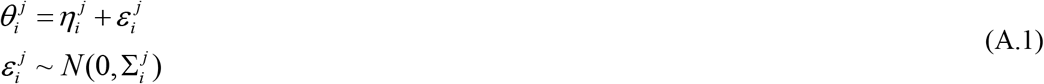

The spectrum is constructed from frequency (*U*) and temporal (*V*) basis functions, which are combined with a matrix of coefficients (*θ^Q^*) corresponding to lexical parameters. The spectrum is scaled with an inverse temperature (i.e., precision; *θ^T^*) parameter, which is then exponentiated to create a matrix of fluctuations *W* of (formant) frequencies over time:

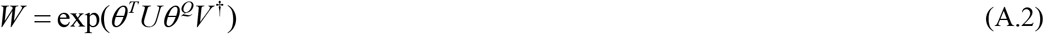

Each column of *W* is transformed into a transient as a function of time (using discrete cosine transform matrix *D*):

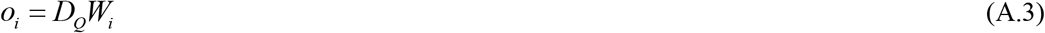

The duration of the transients (*λ*) is determined by the speaker formant spacing (*θ*^1^)—such that a high formant spacing value squashes (shortens) the transients, rendering the frequencies higher when placed in the timeseries. *F_s_* indicates the sampling rate of the audio timeseries:

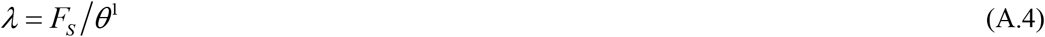

The spacing (Δ) of the transients is inversely proportional to the speaker fundamental frequency parameter (*θ*^0^), and is also affected by inflections due to prosody (*θ^I^*):

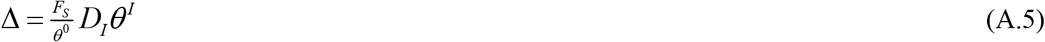

A fundamental interval (*I*) variable stores the absolute positions of all of the transients:

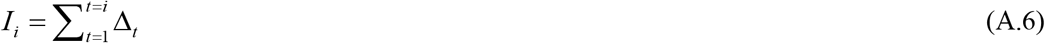

The timeseries (*o*) is constructed by summing the transients and multiplying this by the amplitude

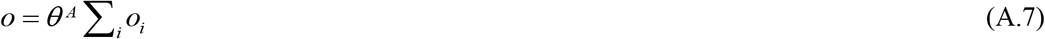

For readers familiar with graphical formulations of generative models, Figure 12 illustrates the same model in factor graph form (Forney 2001). This provides an alternative visual representation of the generative model, and highlights inferences based on message passing. This perspective is used below to describe the form of local (neuronal) message passing that underwrites simulated electrophysiological responses.

**Figure 12.**
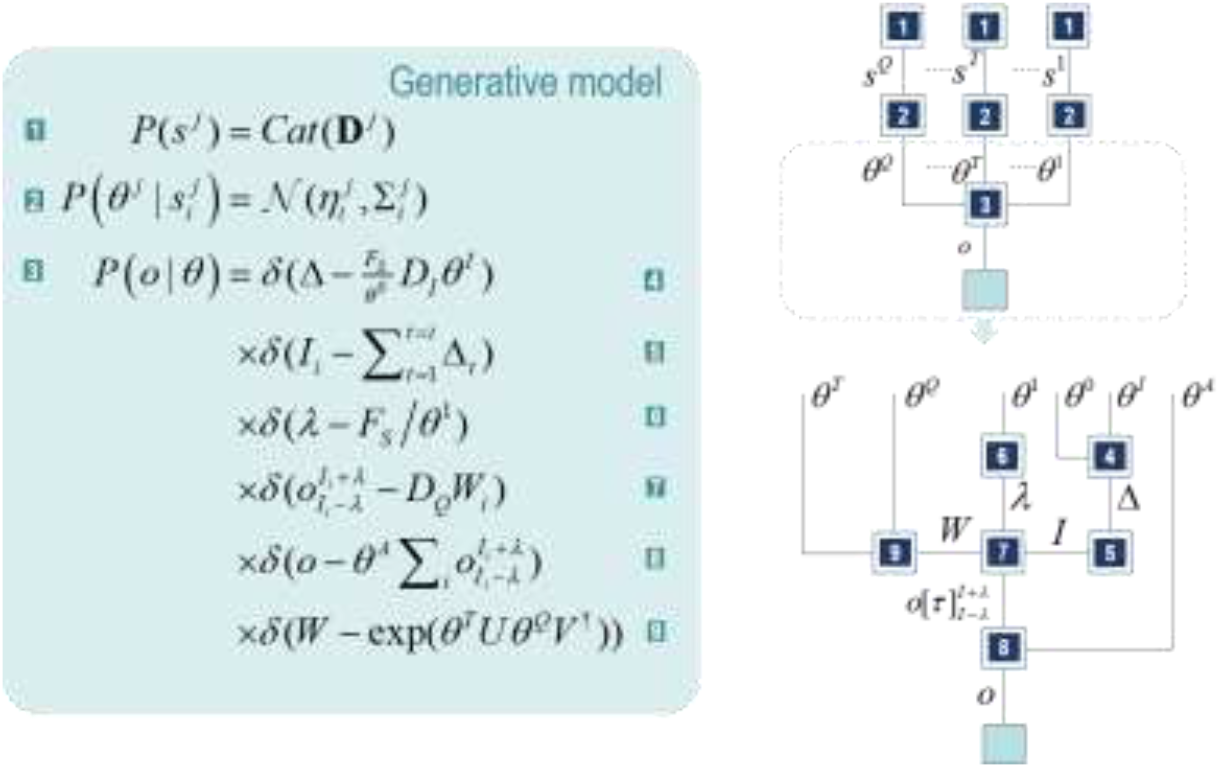
A graphzcal formulatzon of the generatzve model. This figure illustrates the same model as described in Figure 11, but uses a normal (Forney) factor graph form. This graphical notation relies upon the factorisation of the probability density that underwrites the generative model. Each factor is specified in the panel on the left. Factor 1 is the prior probability associated with the hidden states and takes a categorical form. Factor 2 is a normal distribution that specifies the dependence of parameters on states. Each discrete state is associated with a different expectation and covariance for the parameters. Factor 3 describes how the observed timeseries is generated from the parameters, and this is decomposed into factors 4–9. These are Dirac delta functions that may be thought of as normal distributions, centred on zero, with infinite precision (i.e., zero covariance). In the graphs on the right, factors are indicated by numbered squares, and these are connected by edges (Hasson, Yang et al.), which represent the variables common to the factors they connect. The upper right graph shows factors 1–3, and the lower graph unpacks factor 3 in terms of factors 4–9. The process of generating data may be thought of in terms of a series of local operations taking place at each factor from top to bottom (i.e., sample states from factor 1, then parameters from factor 2, then perform the series of operations in factor 3 to get the timeseries). The recognition process can be thought of as bidirectional message passing across each factor node, such that empirical priors and likelihoods are combined at each edge to form posterior beliefs about the associated variable. Factor 5 is of particular interest here, as it determines the internal ‘action’ that selects the interval for segmentation.

## Appendix 2: Model inversion or word recognition

Next, we turn our attention to word recognition (right column of Figure 11). Inversion of the generative model simply requires ‘undoing’ the sequence of events that we used for word generation. Like word generation, word recognition simply requires a series of non-linear transforms—except, for word recognition, we map from epochs of the acoustic signal to discrete *lexical*, *speaker*, and *prosody* parameters.

In brief, the recognition scheme comprises the following steps. The peak energy of the auditory timeseries is identified by convolving its absolute values with a Gaussian kernel. A one second epoch, centred on the peak, is selected as a signal to search for the onset and offset of the word (although in principle this epoch could be any length). Onsets and offsets are identified based on threshold crossings of the amplitude envelope. Here, the amplitude envelope is calculated from the absolute values of the timeseries convolved with a Gaussian kernel. This is, for all practical purposes, equivalent to the absolute values of the Hilbert transform, but is computationally more efficient. The threshold we use here is 1/16^th^ of the maximum envelope value across the window, after subtracting the minimum; this value was selected to be above the noise floor.

The fundamental interval function is estimated using a discrete cosine transform (with three coefficients) of the fundamental intervals. The fundamental intervals are defined as phase crossings following a Hilbert transform and bandpass filtering around the prior for the speaker average fundamental frequency (e.g., 100 Hz, with a standard deviation of 8 Hz).

Equipped with the fundamental interval function, the formant frequencies are then estimated by evaluating the cross-covariance function over short segments centred on each fundamental interval. The duration of these segments corresponds to the inverse of the first formant frequency. The formant frequencies *per se* are evaluated using a modified (by retaining even terms) discrete cosine transform at each slice, to evaluate the spectral density over the acoustic range (in 256 frequency bins, where each bin is determined by the formant spacing; for example, with a formant spacing of 32 Hz, the highest spectral density is 8000 Hz). Following a log transform and normalisation, fluctuations in (log) spectral density are recovered with a discrete cosine transform with 32 basis functions over (formant) frequencies and eight basis functions over intervals. The inverse temperature (timbre) parameter corresponds to the standard deviation of these lexical (formant frequency) parameters, which is used to normalise the lexical (32×8) parameter matrix.

To infer the lexical content, prosody and speaker, the MAP parameter estimates above can be used to evaluate the likelihood of each discrete attribute. As described in the main text, the likelihoods are combined with a prior to produce a posterior categorical distribution over the attributes in question. For the prosody parameters, each parameter is divided into eight bins and the likelihood of belonging to any particular bin is evaluated under Gaussian assumptions as above; using *a priori* means and precisions of the discrete levels of each prosody attribute (i.e., amplitude, duration, timbre, inflection). Similarly, the categorical speaker identity is determined by a 16 x 16 discrete states space, covering fundamental and formant frequencies.

In what follows, we unpack each of the equations in Figure 11—this time, from the perspective of word recognition (right column of Figure 11).

The amplitude parameter is the standard deviation of the timeseries (o):

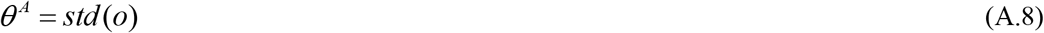

Each transient (*o_i_*) is defined as an interval of the timeseries, based on the positions of fundamental intervals (*I*) and transient durations (*λ*):

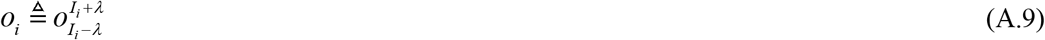

The spacing (Δ) of the transients corresponds to the difference between successive fundamental intervals (*I*):

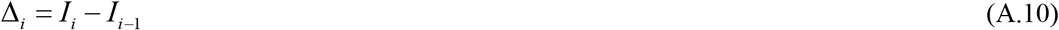

Inflection parameters are proportional to the speaker fundamental frequency (*θ*^0^) and are constructed using discrete cosine transform matrix *D. F_s_* indicates the sampling rate of the audio timeseries:

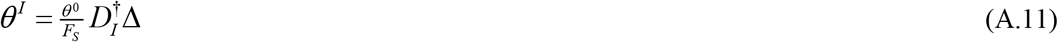

The formant scaling parameter (*θ*^1^) is inversely proportional to the transient duration (*λ*):

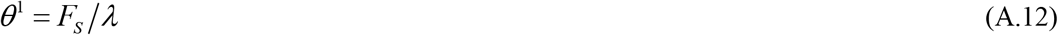

The duration parameter (*θ*^Δ^) is proportional to the fundamental interval (*I*):

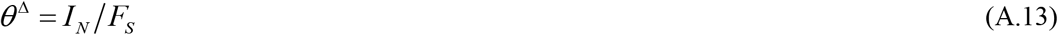

The (squared) matrix of fluctuations of (formant) frequencies over time (*W*) is constructed from the transients using discrete cosine transform matrix *D*:

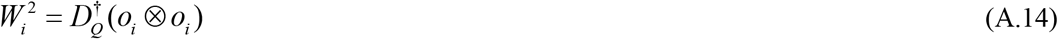

The timbre parameter (*θ^T^*) is the standard deviation of the log spectral decomposition:

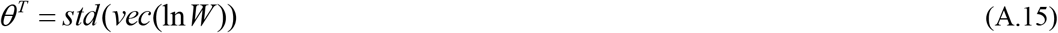

Lexical parameters (*θ^Q^*) are a matrix of coefficients that control the joint expression of formant frequency and temporal basis functions. These are calculated from the frequency (*U*) and temporal (*V*) basis functions and the log spectral decomposition, scaled by the timbre parameter:

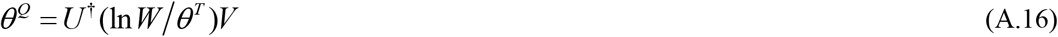

The parameters are used to evaluate the likelihood of lexical, prosody and speaker states, as shown in the following equations:

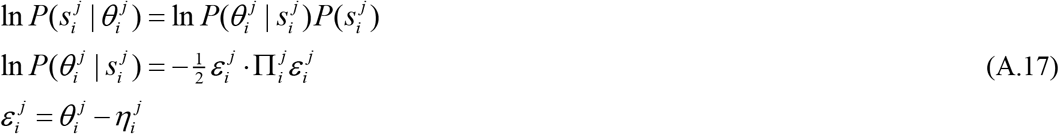

## Appendix 3: Speech segmentation as an active process

In the current framework, speech segmentation is treated as a covert action from a computational perspective: We select boundary pairs (*I*_0_ and *I_T_*) and evaluate their free energy under prior beliefs about the word. Formally, this can be expressed as minimising free energy both with respect to (approximate) posterior beliefs about the attributes of the word (*Q*) and the intervals selected (*I*_0_, *I_T_*):

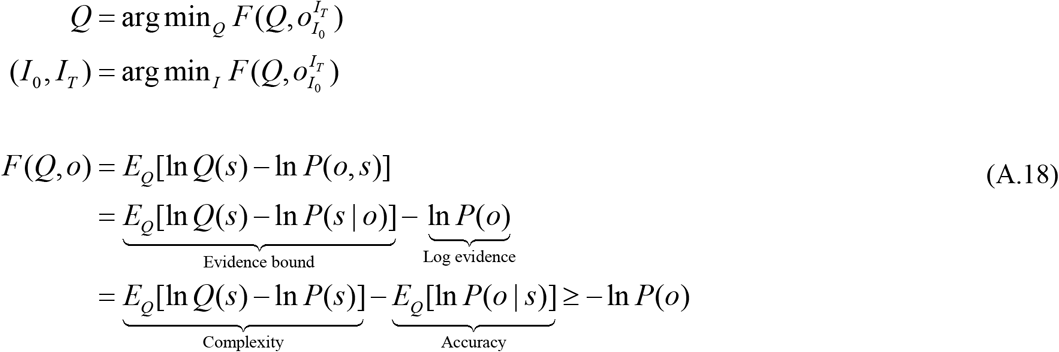

Choosing the interval with the smallest free energy effectively selects the interval that maximises the evidence or marginal likelihood of auditory outcomes contained in that interval; namely, *P*(*o*). This follows because the variational free energy, by construction, represents an upper bound on log evidence. In (A.18), the free energy is expressed in terms of *log evidence* and an evidence *bound*. It is also expressed as the difference between *complexity* and *accuracy* by rearranging the equation. Complexity is the Kullback-Leibler divergence between a posterior over latent states *Q*(*s*), and prior beliefs *P*(*s*), while accuracy is the expected log likelihood of auditory signals contained in the interval in question. Importantly, both posterior beliefs about latent states (i.e., *lexical*, *prosody*, and *speaker*) and the active selection of acoustic intervals optimise free energy. This is the signature of active inference. In this instance, the posterior beliefs obtain from the likelihood of the lexical, prosody and identity parameters, given the associated states. From Figure 11, the optimal posterior beliefs satisfy (A.18) when (ignoring constants):

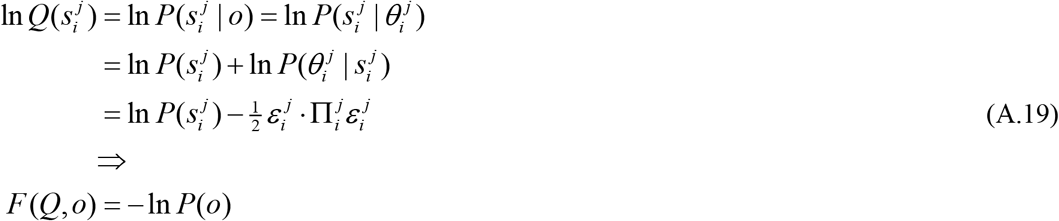

Here, Π is the prior precision of lexical parameters from Figure 11. The second equality on the first line may seem a little counterintuitive, but rests upon the assumed relationship between the parameters and the timeseries. The equality holds in virtue of the absence of random fluctuations in this mapping, such that a given parameter deterministically generates time-series data. In other words, the implicit conditional probability density describing the generation of the timeseries from the parameters (and the associated posterior distribution over parameters) takes the form of a Dirac delta function. The last equality reflects the fact that when the evidence bound in **Error! Reference source not found.** collapses to zero, free energy becomes negative log evidence. The subscript notation indicates the value that a discrete state might take (i.e. *P*(*s*_i_) should be read as ‘the probability that the hidden state *j* takes its *i*-th possible value’).

From the equations above, it should be clear that we can identify a variety of *candidate* boundaries for words and evaluate their free energy to select the final parsing of the acoustic signal. But where should these candidate boundaries be placed? In an extreme case, we could place boundaries at every combination of time points within the acoustic signal—but that would be computationally inefficient given that we can reduce the scope of possibilities by using sensible priors. Here, we use the simple prior that word boundaries are more likely to occur at local minima of the amplitude envelope—so these are the boundaries that we choose to evaluate.

Practically, based upon the spectral content of speech, we estimate the amplitude envelope by removing low frequencies up to about 512 Hz. The envelope is then simply the average of the ensuing absolute values, smoothed with a Gaussian kernel (with a standard deviation of *F_S_*/16). This method is less computationally demanding than using the absolute values of the Hilbert transform, yet practically gives the same result in this setting.

## Appendix 4: Belief updating and neuronal dynamics

The form of neuronal dynamics is calculated by constructing ordinary differential equations whose solution satisfies Equation (A.18). Using **ν** = ln**s** to denote the log of the approximate posterior expectation about hidden states and introducing a prediction error (**ε**) one obtains the following update scheme (Friston, FitzGerald et al. 2017) (dropping the superscript *j* for clarity):

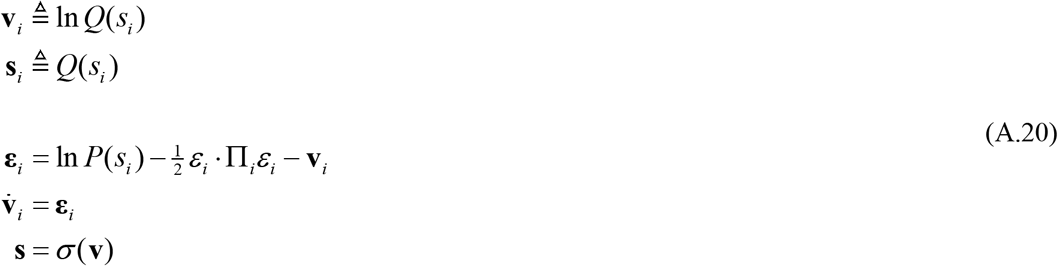

Here, *σ* denotes the softmax (normalised exponential) function and Π is the prior precision of lexical parameters from Figure 11. The prediction error (**ε**) is the difference between the optimal log posterior and current estimate of this (**v**). The log posterior, via Bayes theorem, is equal to the sum of the log prior and the log likelihood (minus a normalisation constant). As the likelihood is assumed to be normally distributed, its log is quadratic in the difference (**ε**) between the mode and lexical parameters. The mode of this distribution is different under each state, so the likelihood of a given parameter value varies with states. For readers familiar with clustering procedures, this is like having a series of clusters (states) with different centroids (i.e., modes of the likelihood).

The prediction error (**ε**) is the (negative) free energy gradient that drives neuronal dynamics. Intuitively, the fourth line of Equation A.20 drives **v** to change until it is equal to the Bayes optimal posterior, at which point **ε** is zero. To account for the normalisation constant that would have appeared in Bayes theorem, the conversion from **v** to **s** requires not only that we exponentiate (i.e., convert a log probability into a probability), but that we normalise the result. This ensures that **s** comes to encode a vector of posterior probabilities for each hidden state.

The sigmoid (softmax) function in Equation A.20 can be thought of as a sigmoid (voltage–firing rate) activation function, which mediates competition among posterior expectations. Equation A.20 therefore, provides a process theory for neuronal dynamics. Based on this equation, log expectations about hidden states can be associated with depolarisation of neurons or neuronal populations encoding expectations about hidden states (**v**_*i*_), while firing rates (**s**_*i*_) encode expectations *per se*. The simulated responses in Figure 6 use a finite difference scheme that has the same solution as A.20:

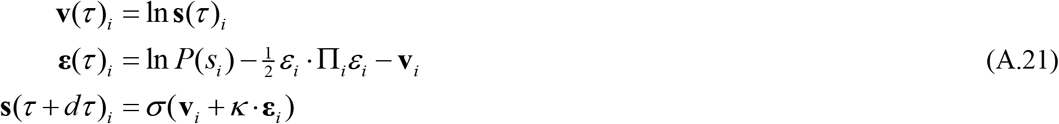

where *κ* is chosen to reproduce dynamics at a plausible, neuronal timescale.

When considering electrophysiological responses in terms of belief updating, our formal interpretation relates to Equation (A.20), which suggests that depolarisation corresponds to the log posterior. The change in depolarisation is the difference between the log posterior and prior expectations. The average of these differences is the Kullback-Leibler divergence between the posterior and prior:

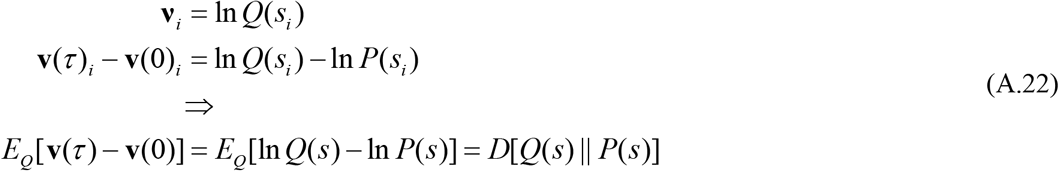

## References

Abberton, E. and A. J. Fourcin (1978). “Intonation and Speaker Identification.” Language and Speech 21(4): 305–318.

Adams, R. A., S. Shipp and K. J. Friston (2013). “Predictions not commands: active inference in the motor system.” Brain Struct Funct. 218(3): 611–643.

Aitchison, L. and M. Lengyel (2017). “With or without you: predictive coding and Bayesian inference in the brain.” Current opinion in neurobiology 46: 219–227.

Alain, C., S. R. Arnott, S. Hevenor, S. Graham and C. L. Grady (2001). ““What” and “where” in the human auditory system.” Proceedings of the National Academy of Sciences 98(21): 12301–12306.

Altenberg, E. P. (2005). “The perception of word boundaries in a second language.” Second Language Research 21(4): 325–358.

Andreopoulos, A. and J. Tsotsos (2013). “A computational learning theory of active object recognition under uncertainty.” International journal of computer vision 101(1): 95–142.

Aring, C. D. (1963). “Traumatic Aphasia: A Study of Aphasia in War Wounds of the Brain.” JAMA Neurology 8(5): 579–580.

Attwell, D. and C. Iadecola (2002). “The neural basis of functional brain imaging signals.” Trends in Neurosciences 25(12): 621–625.

Balaguer, R. D. D., J. M. Toro, A. Rodriguez-Fornells and A.-C. Bachoud-Lévi (2007). “Different neurophysiological mechanisms underlying word and rule extraction from speech.” PLoS One 2(11): e1175.

Bar, M., K. S. Kassam, A. S. Ghuman, J. Boshyan, A. M. Schmid, A. M. Dale, M. S. Hämäläinen, K. Marinkovic, D. L. Schacter and B. R. Rosen (2006). “Top-down facilitation of visual recognition.” Proceedings of the national academy of sciences 103(2): 449–454.

Barto, A., M. Mirolli and G. Baldassarre (2013). “Novelty or Surprise?” Frontiers in Psychology 4.

Bashford, J. A., Jr., R. M. Warren and P. W. Lenz (2008). “Evoking biphone neighborhoods with verbal transformations: illusory changes demonstrate both lexical competition and inhibition.” J Acoust Soc Am 123(3): E132.

Bastos, A. M., W. M. Usrey, R. A. Adams, G. R. Mangun, P. Fries and K. J. Friston (2012). “Canonical microcircuits for predictive coding.” Neuron 76(4): 695–711.

Baumann, O. and P. Belin (2009). “Perceptual scaling of voice identity: Common dimensions for different vowels and speakers.” Psychological Research 74(1): 110–120.

Beal, M. J. (2003). “Variational Algorithms for Approximate Bayesian Inference.” PhD. Thesis, University College London.

Beckman, M. E. and J. Edwards (1990). “of prosodic constituency.” Between the grammar and physics of speech: 152.

Belin, P., S. Fecteau and C. Bdard (2004). “Thinking the voice: Neural correlates of voice perception.” Trends in Cognitive Sciences 8(3): 129–-135.

Belin, P. and R. J. Zatorre (2000). “‘What’, ‘where’ and ‘how’ in auditory cortex.” Nature Neuroscience 3(10): 965–966.

Bennett, C. H. (2003). “Notes on Landauer’s principle, reversible computation, and Maxwell’s Demon.” Studies in History and Philosophy of Science Part B: Studies in History and Philosophy of Modern Physics 34(3): 501–510.

Billig, A. J., M. H. Davis, J. M. Deeks, J. Monstrey and R. P. Carlyon (2013). “Lexical influences on auditory streaming.” Current Biology 23(16): 1585–-1589.

Bogacz, R. (2017). “A tutorial on the free-energy framework for modelling perception and learning.” Journal of Mathematical Psychology 76: 198–-211.

Bögels, S., H. Schriefers, W. Vonk and D. J. Chwilla (2011). “Prosodic Breaks in Sentence Processing Investigated by Event-Related Potentials.” Language and Linguistics Compass 5(7): 424–440.

Bögels, S., H. Schriefers, W. Vonk and D. J. Chwilla (2011). “The role of prosodic breaks and pitch accents in grouping words during on-line sentence processing.” Journal of Cognitive Neuroscience 23(9): 2447–2467.

Braiman, C., E. A. Fridman, M. M. Conte, H. U. Voss, C. S. Reichenbach, T. Reichenbach and N. D. Schiff (2018). “Cortical Response to the Natural Speech Envelope Correlates with Neuroimaging Evidence of Cognition in Severe Brain Injury.” Curr Biol 28(23): 3833–3839.e3833.

Brown, H., R. A. Adams, I. Parees, M. Edwards and K. J. Friston (2013). “Active inference, sensory attenuation and illusions.” Cognitive Processing 14(4): 411–-427.

Brown, H., K. J. Friston and S. Bestmann (2011). “Active inference, attention, and motor preparation.” Frontiers in psychology 2: 218.

Brungart, D. S. (2001). “Evaluation of speech intelligibility with the coordinate response measure.” The Journal of the Acoustical Society of America 109(5 Pt 1): 2276–-2279.

Brungart, D. S., B. D. Simpson, M. A. Ericson and K. R. Scott (2001). “Informational and energetic masking effects in the perception of multiple simultaneous talkers.” The Journal of the Acoustical Society of America 110(5): 2527–2538.

Cai, Z. G., R. A. Gilbert, M. H. Davis, M. G. Gaskell, L. Farrar, S. Adler and J. M. Rodd (2017). “Accent modulates access to word meaning: Evidence for a speaker-model account of spoken word recognition.” Cognitive Psychology 98: 73–101.

Christie Jr, W. M. (1974). “Some cues for syllable juncture perception in English.” the Journal of the Acoustical Society of America 55(4): 819–821.

Cole, R. A., J. Jakimik and W. E. Cooper (1980). “Segmenting speech into words.” The Journal of the Acoustical Society of America 67(4): 1323–1332.

Connolly, J. F. and N. A. Phillips (1994). “Event-related potential components reflect phonological and semantic processing of the terminal word of spoken sentences.” Journal of cognitive neuroscience 6(3): 256–266.

Connolly, J. F., N. A. Phillips, S. H. Stewart and W. Brake (1992). “Event-related potential sensitivity to acoustic and semantic properties of terminal words in sentences.” Brain and language 43(1): 1–18.

Cunillera, T., E. Càmara, J. M. Toro, J. Marco-Pallares, N. Sebastián-Galles, H. Ortiz, J. Pujol and A. Rodríguez-Fornells (2009). “Time course and functional neuroanatomy of speech segmentation in adults.” Neuroimage 48(3): 541–553.

Cutler, A. and D. Norris (1988). “The role of strong syllables in segmentation for lexical access.” Journal of Experimental Psychology: Human perception and performance 14(1): 113.

Davis, M. H. and I. S. Johnsrude (2003). “Hierarchical processing in spoken language comprehension.” Journal of Neuroscience 23(8): 3423–3431.

Davis, M. H., W. D. Marslen-Wilson and M. G. Gaskell (2002). “Leading up the lexical garden path: Segmentation and ambiguity in spoken word recognition.” Journal of Experimental Psychology: Human Perception and Performance 28(1): 218.

Davison, A. J. and D. W. Murray (2002). “Simultaneous localization and map-building using active vision.” Ieee Transactions on Pattern Analysis and Machine Intelligence 24(7): 865–880.

Dehaene-Lambertz, G. (1997). “Electrophysiological correlates of categorical phoneme perception in adults.” NeuroReport 8(4): 919–924.

DeWitt, I. and J. P. Rauschecker (2012). “Phoneme and word recognition in the auditory ventral stream.” Proceedings of the National Academy of Sciences of the United States of America 109(8): E505–E514.

Ding, N., L. Melloni, H. Zhang, X. Tian and D. Poeppel (2015). “Cortical tracking of hierarchical linguistic structures in connected speech.” Nature Neuroscience 19(1): 158–-164.

Ding, N. and J. Z. Simon (2012). “Neural coding of continuous speech in auditory cortex during monaural and dichotic listening.” Journal of neurophysiology 107(1): 78–-89.

Donchin, E. and M. G. H. Coles (1988). “Is the P300 component a manifestation of context updating?” Behavioral and Brain Sciences 11(3): 357.

Drullman, R. (1995). “Temporal envelope and fine structure cues for speech intelligibility.” Journal of the Acoustical Society of America 97(1): 585–592.

Dubno, J. R., J. B. Ahlstrom and a. R. Horwitz (2000). “Use of context by young and aged adults with normal hearing.” The Journal of the Acoustical Society of America 107(1): 538–-546.

Durlach, N. (2006). “Auditory masking: Need for improved conceptual structure.” The Journal of the Acoustical Society of America 120(4): 1787–1790.

Durlach, N. I., C. R. Mason, G. K. Jr., T. L. Arbogast, H. S. Colburn and B. G. Shinn-Cunningham (2003). “Note on informational masking (L).” The Journal of the Acoustical Society of America 113(6): 2984–2987.

Durlach, N. I., C. R. Mason, B. G. Shinn-Cunningham, T. L. Arbogast, H. S. Colburn and G. Kidd (2003). “Informational masking: Counteracting the effects of stimulus uncertainty by decreasing target-masker similarity.” The Journal of the Acoustical Society of America 114(1): 368.

Easwar, V., D. W. Purcell, S. J. Aiken, V. Parsa and S. D. Scollie (2015). “Evaluation of Speech-Evoked Envelope Following Responses as an Objective Aided Outcome Measure: Effect of Stimulus Level, Bandwidth, and Amplification in Adults With Hearing Loss.” Ear Hear 36(6): 635–652.

Feldman, A. G. and M. F. Levin (1995). “The origin and use of positional frames of reference in motor control.” Behav Brain Sci. 18: 723–806.

Feynman, R. P. (1972). Statistical mechanics. Reading MA, Benjamin.

Fiez, J. A. and S. E. Petersen (1998). “Neuroimaging studies of word reading.” Proc Natl Acad Sci U S A 95(3): 914–921.

Fitch, W. T. (1997). “Vocal tract length and formant frequency dispersion correlate with body size in rhesus macaques.” The Journal of the Acoustical Society of America 102(2): 1213–1222.

Forney, G. D. (2001). “Codes on graphs: Normal realizations.” IEEE Transactions on Information Theory 47(2): 520548.

François, C., T. Cunillera, E. Garcia, M. Laine and A. Rodriguez-Fornells (2017). “Neurophysiological evidence for the interplay of speech segmentation and word-referent mapping during novel word learning.” Neuropsycholo gia 98: 56–67.

Friederici, A. D., A. Hahne and A. Mecklinger (1996). “Temporal structure of syntactic parsing: early and late event-related brain potential effects.” Journal of Experimental Psychology: Learning, Memory, and Cognition 22(5): 1219.

Friston, K. (2013). “Life as we know it.” J R Soc Interface 10(86): 20130475.

Friston, K. and G. Buzsaki (2016). “The Functional Anatomy of Time: What and When in the Brain.” Trends Cogn Sci.

Friston, K., T. FitzGerald, F. Rigoli, P. Schwartenbeck and G. Pezzulo (2017). “Active Inference: A Process Theory.” Neural Comput 29(1): 1–49.

Friston, K. and C. Frith (2015). “A duet for one.” Consciousness and cognition 36: 390–405.

Friston, K., J. Mattout and J. Kilner (2011). “Action understanding and active inference.” Biol Cybern. 104: 137–160.

Friston, K. J. (2010). “The free-energy principle: A unified brain theory?” Nature Reviews Neuroscience 11(2): 127–138.

Friston, K. J., T. FitzGerald, F. Rigoli, P. Schwartenbeck and G. Pezzulo (2017). “Active Inference: A Process Theory.” Neural computation 29: 1–-49.

Friston, K. J., T. Parr and B. de Vries (2017). “The graphical brain: belief propagation and active inference.” Network Neuroscience: 1–-78.

Friston, K. J., T. Parr and B. de Vries (2017). “The graphical brain: Belief propagation and active inference.” Netw Neurosci 1(4): 381–414.

Friston, K. J., R. Rosch, T. Parr, C. Price and H. Bowman (2017). “Deep temporal models and active inference.” Neurosci Biobehav Rev 77: 388–402.

Ganong, W. F. (1980). “Phonetic categorization in auditory word perception.” Journal of experimental psychology: Human perception and performance 6(1): 110.

Garrido, M. I., J. M. Kilner, K. E. Stephan and K. J. Friston (2009). “The mismatch negativity: a review of underlying mechanisms.” Clin Neurophysiol 120(3): 453–463.

Gaskell, M. G. and W. D. Marslen-Wilson (1997). “Integrating form and meaning: A distributed model of speech perception.” Language and cognitive Processes 12(5-6): 613–656.

Gaudrain, E., S. Li, V. S. Ban and R. D. Patterson (2009). “The role of glottal pulse rate and vocal tract length in the perception of speaker identity.” Proceedings of the Annual Conference of the International Speech Communication Association, INTERSPEECH(January 2009): 148–-151.

Giard, M., J. Lavikahen, K. Reinikainen, F. Perrin, O. Bertrand, J. Pernier and R. Näätänen (1995). “Separate representation of stimulus frequency, intensity, and duration in auditory sensory memory: an event-related potential and dipole-model analysis.” Journal of cognitive neuroscience 7(2): 133–143.

Gow Jr, D. W. and P. C. Gordon (1995). “Lexical and prelexical influences on word segmentation: Evidence from priming.” Journal of Experimental Psychology: Human perception and performance 21(2): 344.

Grossberg, S., K. Roberts, M. Aguilar and D. Bullock (1997). “A neural model of multimodal adaptive saccadic eye movement control by superior colliculus.” J Neurosci. 17(24): 9706–9725.

Grotheer, M. and G. Kovács (2014). “Repetition probability effects depend on prior experiences.” The Journal of neuroscience: the official journal of the Society for Neuroscience 34 19: 6640–6646.

Guenther, F. H. and T. Vladusich (2012). “A Neural Theory of Speech Acquisition and Production.” J Neurolinguistics 25(5): 408–422.

Harris, M. and N. Umeda (1974). “Effect of speaking mode on temporal factors in speech: Vowel duration.” The Journal of the Acoustical Society of America 56(3): 1016–1018.

Hasson, U., E. Yang, I. Vallines, D. J. Heeger and N. Rubin (2008). “A hierarchy of temporal receptive windows in £1 human cortex.” J Neurosci 28(10): 2539–2550.

Heilbron, M. and M. Chait (2018). “Great Expectations: Is there Evidence for Predictive Coding in Auditory Cortex?” Neuroscience 389: 54–73.

Hickok, G. (2014). “The architecture of speech production and the role of the phoneme in speech processing.” Lang Cogn Process 29(1): 2–20.

Hickok, G. and D. Poeppel (2007). “Opinion - The cortical organization of speech processing.” Nature Reviews Neuroscience 8(5): 393–402.

Hillenbrand, J. M., L. A. Getty, M. J. Clark and K. Wheeler (1995). “Acoustic characteristics of American English vowels.” Journal of the Acoustical Society of America 97(5): 3099–-3111.

Hinton, G. E. and R. S. Zemel (1993). Autoencoders, minimum description length and Helmholtz free energy. Proceedings of the 6th International Conference on Neural Information Processing Systems. Denver, Colorado, Morgan Kaufmann Publishers Inc.:3–10.

Hodges, J. R., K. Patterson, S. Oxbury and E. Funnell (1992). “Semantic dementia. Progressive fluent aphasia with temporal lobe atrophy.” Brain 115 (Pt 6): 1783–1806.

Hohwy, J. (2016). “The Self-Evidencing Brain.” Noûs 50(2): 259–285.

Holmes, E., Y. Domingo and I. S. Johnsrude (2018). “Familiar voices are more intelligible, even if they are not recognized as familiar.” Psychological Science 29(10): 1575–-1583.

Holmes, E., P. Folkeard, I. S. Johnsrude and S. Scollie (2018). “Semantic context improves speech intelligibility and reduces listening effort for listeners with hearing impairment.” Int J Audiol 57(7): 483–492.

Holt, L. L., A. J. Lotto and K. R. Kluender (2000). “Neighboring spectral content influences vowel identification.” Journal of the Acoustical Society of America 108(2): 710–722.

Hope, T. M. H., A. P. Leff and C. J. Price (2018). “Predicting language outcomes after stroke: Is structural disconnection a useful predictor?” NeuroImage. Clinical 19: 22–29.

Houde, J. and S. Nagarajan (2011). “Speech Production as State Feedback Control.” Frontiers in Human Neuroscience 5(82).

Itti, L. and P. Baldi (2009). “Bayesian Surprise Attracts Human Attention.” Vision Res. 49(10): 1295–1306.

Jacobsen, T., T. Horenkamp and E. Schröger (2003). “Preattentive memory-based comparison of sound intensity.” Audiology and Neurotology 8(6): 338–346.

Jacobsen, T., E. Schröger, T. Horenkamp and I. Winkler (2003). “Mismatch negativity to pitch change: varied stimulus proportions in controlling effects of neural refractoriness on human auditory event-related brain potentials.” Neuroscience letters 344(2): 79–82.

Johnsrude, I. S., A. Mackey, H. Hakyemez, E. Alexander, H. P. Trang and R. P. Carlyon (2013). “Swinging at a cocktail party: voice familiarity aids speech perception in the presence of a competing voice.” Psychological science 24(10): 1995–-2004.

Kaas, J. H. and T. A. Hackett (1999). “‘What’ and ‘where’ processing in auditory cortex.” Nat Neurosci 2(12): 1045–1047.

Kidd, G., C. R. Mason, V. M. Richards, F. Gallun and N. Durlach (2007). Informational Masking. 29:143–189.

Kiebel, S. J., J. Daunizeau and K. J. Friston (2009). “Perception and hierarchical dynamics.” Front Neuroinform 3: 20.

Kim, D., J. D. Stephens and M. A. Pitt (2012). “How does context play a part in splitting words apart? Production and perception of word boundaries in casual speech.” Journal of memory and language 66(4): 509–529.

Kim, S., R. D. Frisina, F. M. Mapes, E. D. Hickman and D. R. Frisina (2006). “Effect of age on binaural speech intelligibility in normal hearing adults.” Speech Communication 48(6): 591–-597.

Klatt, D. H. (1975). “Vowel lengthening is syntactically determined in a connected discourse.” Journal of phonetics 3(3): 129–140.

Klatt, D. H. (1976). “Linguistic uses of segmental duration in English: Acoustic and perceptual evidence.” The Journal of the Acoustical Society of America 59(5): 1208–1221.

Kleinschmidt, D. F. and T. F. Jaeger (2015). “Robust Speech Perception: Recognize the Familiar, Generalize to the Similar, and Adapt to the Novel.” Psychological Review 122(2): 148–203.

Kuhlen, A. K., C. Bogler, S. E. Brennan and J.-D. Haynes (2017). “Brains in dialogue: decoding neural preparation of speaking to a conversational partner.” Social cognitive and affective neuroscience 12(6): 871–880.

Kumar, S., K. E. Stephan, J. D. Warren, K. J. Friston and T. D. Griffiths (2007). “Hierarchical processing of auditory objects in humans.” PLoS computational biology 3(6): e100.

Kuperberg, G. R., T. Sitnikova, D. Caplan and P. J. Holcomb (2003). “Electrophysiological distinctions in processing conceptual relationships within simple sentences.” Cognitive brain research 17(1): 117–129.

Kutas, M. and K. D. Federmeier (2000). “Electrophysiology reveals semantic memory use in language comprehension.” Trends in cognitive sciences 4(12): 463–470.

Kutas, M. and K. D. Federmeier (2009). “N400.” Scholarpedia 4(10): 7790.

Kutas, M. and S. A. Hillyard (1980). “Reading senseless sentences: Brain potentials reflect semantic incongruity.” Science 207(4427): 203–205.

Kutas, M. and S. A. Hillyard (1984). “Brain potentials during reading reflect word expectancy and semantic association.” Nature 307(5947): 161.

Ladd, D. R. and A. Schepman (2003). ““Sagging transitions” between high pitch accents in English: Experimental evidence.” Journal of phonetics 31(1): 81–112.

Landauer, R. (1961). “Irreversibility and Heat Generation in the Computing Process.” IBM Journal of Research and Development 5(3): 183–191.

Landi, N., S. J. Frost, W. E. Menc, R. Sandak and K. R. Pugh (2013). “Neurobiological bases of reading comprehension: Insights from neuroimaging studies of word level and text level processing in skilled and impaired readers.” Read Writ Q 29(2): 145–167.

LaRiviere, C. (1975). “Contributions of Fundamental Frequency and Formant Frequencies to Speaker Identification.” Phonetica 31(3-4): 185–197.

Larsson, J. and A. T. Smith (2012). “fMRI repetition suppression: neuronal adaptation or stimulus expectation?” Cereb Cortex 22(3): 567–576.

Lavner, Y., I. Gath and J. Rosenhouse (2000). “Effects of acoustic modifications on the identification of familiar voices speaking isolated vowels.” Speech Communication 30(1): 9–-26.

Lavner, Y., J. Rosenhouse and I. Gath (2001). “The prototype model in speaker identification by human listeners.” International Journal of Speech Technology 4(1): 63–-74.

Lehiste, I. (1960). “An acoustic-phonetic study of internal open juncture.” Phonetica 5(Suppl. 1): 5–54.

Lehiste, I. (1972). “The timing of utterances and linguistic boundaries.” The Journal of the Acoustical Society of America 51(6B): 2018–2024.

Lehiste, I. (1973). “Rhythmic units and syntactic units in production and perception.” The Journal of the Acoustical Society of America 54(5): 1228–1234.

Liberman, A. M., F. S. Cooper, D. P. Shankweiler and M. Studdert-Kennedy (1967). “Perception of the speech code.” Psychological review 74(6): 431.

Luce, P. A. (1986). “Neighborhoods of words in the mental lexicon.” Research on speech perception, Technical Report 6: 1–91.

Luce, P. A. and D. B. Pisoni (1998). “Recognizing spoken words: the neighborhood activation model.” Ear and hearing 19(1): 1–36.

Maisto, D., F. Donnarumma and G. Pezzulo (2015). “Divide et impera: subgoaling reduces the complexity of probabilistic inference and problem solving.” 12(104): 20141335.

Mann, V. A. (1980). “Influence of preceding liquid on stop-consonant perception.” Perception & Psychophysics 28(5): 407–412.

Marslen-Wilson, W. D. (1975). “Sentence perception as an interactive parallel process.” Science 189(4198): 226–228.

Marslen-Wilson, W. D. (1984). Function and process in spoken word recognition: A tutorial review. Attention and performance: Control of language processes, Erlbaum:125–150.

Marslen-Wilson, W. D. and A. Welsh (1978). “Processing interactions and lexical access during word recognition in continuous speech.” Cognitive psychology 10(1): 29–63.

Massaro, D. W. (1987). Categorical partition: A fuzzy-logical model of categorization behavior. Categorical perception: The groundwork of cognition. New York, NY, US, Cambridge University Press:254–283.

Massaro, D. W. (1989). “Testing between the TRACE model and the fuzzy logical model of speech perception.” Cognitive psychology 21(3): 398–421.

Matsumoto, H., S. Hiki, T. Sone and T. Nimura (1973). “Multidimensional representation of personal quality of vowels and its acoustical correlates.” IEEE Transactions on Audio and Electroacoustics 21(5): 428–-436.

Mattys, S. L. and J. F. Melhorn (2007). “Sentential, lexical, and acoustic effects on the perception of word boundaries.” The Journal of the Acoustical Society of America 122(1): 554–567.

Mattys, S. L., J. F. Melhorn and L. White (2007). “Effects of syntactic expectations on speech segmentation.” Journal of Experimental Psychology: Human Perception and Performance 33(4): 960.

Mattys, S. L., L. White and J. F. Melhorn (2005). “Integration of multiple speech segmentation cues: A hierarchical framework.” Journal of Experimental Psychology-General 134(4): 477–500.

McClelland, J. L. and J. L. Elman (1986). “The TRACE model of speech perception.” Cognitive Psychology 18(1): 1–86.

Mermelstein, P. (1967). “Determination of the Vocal-Tract Shape from Measured Formant Frequencies.” The Journal of the Acoustical Society of America 41(5): 1283–1294.

Miller, J. L., K. Green and T. M. Schermer (1984). “A distinction between the effects of sentential speaking rate and semantic congruity on word identification.” Perception & Psychophysics 36(4): 329–337.

Miller, J. L. and A. M. Liberman (1979). “Some effects of later-occurring information on the perception of stop consonant and semivowel.” Perception & Psychophysics 25(6): 457–465.

Mirza, M. B., R. A. Adams, C. D. Mathys and K. J. Friston (2016). “Scene Construction, Visual Foraging, and Active Inference.” Frontiers in Computational Neuroscience 10(56).

Mirza, M. B., R. A. Adams, C. D. Mathys and K. J. Friston (2016). “Scene Construction, Visual Foraging, and Active Inference.” Front Comput Neurosci 10: 56.

Mohan, V. and P. Morasso (2011). “Passive motion paradigm: an alternative to optimal control.” Front Neurorobot 5: 4.

Morlet, D. and C. Fischer (2014). “MMN and novelty P3 in coma and other altered states of consciousness: a review.” Brain Topogr 27(4): 467–479.

Muralimanohar, R. K., J. M. Kates and K. H. Arehart (2017). “Using envelope modulation to explain speech intelligibility in the presence of a single reflection.” J Acoust Soc Am 141(5): El482.

Murry, T. and S. Singh (1980). “Multidimensional analysis of male and female voices.” The Journal of the Acoustical Society of America 68(5): 1294–-1300.

Musso, M., C. Weiller, A. Horn, V. Glauche, R. Umarova, J. Hennig, A. Schneider and M. Rijntjes (2015). “A single dual-stream framework for syntactic computations in music and language.” Neuroimage 117: 267–283.

Näätänen, R., A. W. Gaillard and S. Mäntysalo (1978). “Early selective-attention effect on evoked potential reinterpreted.” Acta psychologica 42(4): 313–329.

Näätänen, R., A. Lehtokoski, M. Lennes, M. Cheour, M. Huotilainen, A. Iivonen, M. Vainio, P. Alku, R. J. Ilmoniemi and A. Luuk (1997). “Language-specific phoneme representations revealed by electric and magnetic brain responses.” Nature 385(6615): 432.

Nakatani, L. H. and K. D. Dukes (1977). “Locus of segmental cues for word juncture.” The Journal of the Acoustical Society of America 62(3): 714–719.

Nardo, D., R. Holland, A. P. Leff, C. J. Price and J. T. Crinion (2017). “Less is more: neural mechanisms underlying anomia treatment in chronic aphasic patients.” Brain 140(11): 3039–3054.

Nealey, T. A. and J. H. Maunsell (1994). “Magnocellular and parvocellular contributions to the responses of neurons in macaque striate cortex.” The Journal of Neuroscience 14(4): 2069.

Norris, D. and J. M. McQueen (2008). “Shortlist B: A Bayesian model of continuous speech recognition.” Psychological review 115(2): 357–-395.

Norris, D., J. M. McQueen and A. Cutler (2016). “Prediction, Bayesian inference and feedback in speech recognition.” Lang Cogn Neurosci 31(1): 4–18.

Norris, D., J. M. McQueen, A. Cutler and S. Butterfield (1997). “The possible-word constraint in the segmentation of continuous speech.” Cognitive Psychology 34(3): 191–243.

Nygaard, L. C., M. S. Sommers and D. B. Pisoni (1994). “SPEECH PERCEPTION AS A TALKER-CONTINGENT PROCESS.” Psychol Sci 5(1): 42–46.

O’Leary, D. D. M. (1989). “Do cortical areas emerge from a protocortex?” Trends in Neurosciences 12(10): 400–406.

O’Sullivan, J. A., A. J. Power, N. Mesgarani, S. Rajaram, J. J. Foxe, B. G. Shinn-Cunningham, M. Slaney, S. a. Shamma and E. Lalor (2014). “Attentional selection in a cocktail party environment can be decoded from single-trial EEG.” Cerebral Cortex: 1–-10.

Oden, G. C. and D. W. Massaro (1978). “Integration of featural information in speech perception.” Psychological review 85(3): 172.

Ognibene, D. and G. Baldassarre (2014). Ecological Active Vision: Four Bio-Inspired Principles to Integrate Bottom-Up and Adaptive Top-Down Attention Tested With a Simple Camera-Arm Robot. IEEE Transactions onAutonomous Mental Development, IEEE.

Oller, D. K. (1973). “The effect of position in utterance on speech segment duration in English.” The journal of the Acoustical Society of America 54(5): 1235–1247.

Osterhout, L. and P. J. Holcomb (1992). “Event-related brain potentials elicited by syntactic anomaly.” Journal of memory and language 31(6): 785–806.

Oudeyer, P.-Y. and F. Kaplan (2007). “What is intrinsic motivation? a typology of computational approaches.” Frontiers in Neurorobotics 1: 6.

Pannekamp, A., U. Toepel, K. Alter, A. Hahne and A. D. Friederici (2005). “Prosody-driven sentence processing: An event-related brain potential study.” Journal of cognitive neuroscience 17(3): 407–421.

Parr, T. and K. J. Friston (2017). “The active construction of the visual world.” Neuropsychologia 104: 92–101.

Parr, T. and K. J. Friston (2017). “Working memory, attention, and salience in active inference.” Scientific Reports 7(1): 14678.

Parr, T., D. Markovic, S. J. Kiebel and K. J. Friston (2019). “Neuronal message passing using Mean-field, Bethe, and Marginal approximations.” Scientific Reports 9(1): 1889.

Pasley, B. N., S. V. David, N. Mesgarani, A. Flinker, S. A. Shamma, N. E. Crone, R. T. Knight and E. F. Chang (2012). “Reconstructing speech from human auditory cortex.” PLoS biology 10(1): e1001251.

Patel, A. D. (2010). Music, language, and the brain. Oxford, UK, Oxford Univ. Press.

Paulesu, E., B. Goldacre, P. Scifo, S. F. Cappa, M. C. Gilardi, I. Castiglioni, D. Perani and F. Fazio (1997). “Functional heterogeneity of left inferior frontal cortex as revealed by fMRI.” Neuroreport 8(8): 2011–2017.

Pearce, M. T. (2018). “Statistical learning and probabilistic prediction in music cognition: mechanisms of stylistic enculturation.” Ann N Y Acad Sci.

Penny, W. D. (2012). “Comparing dynamic causal models using AIC, BIC and free energy.” Neuroimage 59(1): 319330.

Peretz, I., R. Kolinsky, M. Tramo, R. Labrecque, C. Hublet, G. Demeurisse and S. Belleville (1994). “Functional dissociations following bilateral lesions of auditory cortex.” Brain 117(6): 1283–1301.

Picton, T. W., C. Alain, L. Otten, W. Ritter and A. Achim (2000). “Mismatch negativity: different water in the same river.” Audiology and Neurotology 5(3-4): 111–139.

Poeppel, D. and P. J. Monahan (2011). “Feedforward and feedback in speech perception: Revisiting analysis by synthesis.” Language and Cognitive Processes 26(7): 935–951.

Polich, J. (2007). “Updating P300: an integrative theory of P3a and P3b.” Clinical neurophysiology 118(10): 21282148.

Polich, J. and E. Donchin (1988). “P300 and the word frequency effect.” Electroencephalography and clinical neurophysiology 70(1): 33–45.

Price, C. J. (2012). “A review and synthesis of the first 20 years of PET and fMRI studies of heard speech, spoken language and reading.” NeuroImage 62(2): 816–847.

Quiroga-Martinez, D. R., N. C. Hansen, A. Højlund, M. Pearce, E. Brattico and P. Vuust (2019). “Reduced prediction error responses in high-as compared to low-uncertainty musical contexts.” bioRxiv: 422949.

Remez, R. E. (2010). “Spoken expression of individual identity and the listener.” Expressing oneself/expressing one’s self: Communication, cognition, language, and identity.: 167–-181.

Romanski, L. M., B. Tian, J. Fritz, M. Mishkin, P. S. Goldman-Rakic and J. P. Rauschecker (1999). “Dual streams of auditory afferents target multiple domains in the primate prefrontal cortex.” Nat Neurosci 2(12): 1131–1136.

Rosenfeld, R. (2000). “Two decades of statistical language modeling: Where do we go from here?” Proceedings of the Ieee 88(8): 1270–1278.

Rueschemeyer, S.-A., M. G. Gaskell, G. Walker and G. Hickok (2018). Speech ProductionIntegrating psycholinguistic, neuroscience, and motor control perspectives, Oxford University Press.

Ryan, R. and E. Deci (1985). Intrinsic motivation and self-determination in human behavior. New York, Plenum.

Sams, M., P. Paavilainen and K. Alho (1985). “Auditory frequency discrimination and event-related potentials.” Electroencephalography and Clinical Neurophysiology 62: 437–-448.

Sato, Y., H. Yabe, T. Hiruma, T. Sutoh, N. Shinozaki, T. Nashida and S. Kaneko (2000). “The effect of deviant stimulus probability on the human mismatch process.” Neuroreport 11(17): 3703–3708.

Sato, Y., H. Yabe, J. Todd, P. Michie, N. Shinozaki, T. Sutoh, T. Hiruma, T. Nashida, T. Matsuoka and S. Kaneko (2003). “Impairment in activation of a frontal attention-switch mechanism in schizophrenic patients.” Biological psychology 62(1): 49–63.

Schmidhuber, J. (1991). “Curious model-building control systems.” In Proc. International Joint Conference on Neural Networks, Singapore. IEEE 2: 1458–1463.

Schmidhuber, J. (2006). “Developmental robotics, optimal artificial curiosity, creativity, music, and the fine arts.” Connection Science 18(2): 173–187.

Sengupta, B., M. B. Stemmler and K. J. Friston (2013). “Information and efficiency in the nervous system—a synthesis.” PLoS computational biology 9(7): e1003157.

Sengupta, B., A. Tozzi, G. K. Cooray, P. K. Douglas and K. J. Friston (2016). “Towards a Neuronal Gauge Theory.” PLoS Biol 14(3): e1002400.

Seth, A. (2014). The cybernetic brain: from interoceptive inference to sensorimotor contingencies. MINDS project. Metzinger, T; Windt, JM, MINDS.

Shamma, S. (2001). “On the role of space and time in auditory processing.” Trends in cognitive sciences 5(8): 340348.

Shamma, S. A., M. Elhilali and C. Micheyl (2011). “Temporal coherence and attention in auditory scene analysis.” Trends in neurosciences 34(3): 114–-123.

Shiell, M. M., F. Champoux and R. J. Zatorre (2015). “Reorganization of auditory cortex in early-deaf people: Functional connectivity and relationship to hearing aid use.” Journal of Cognitive Neuroscience 27(1): 150–163.

Shillcock, R. (1990). “Lexical hypotheses in continuous speech.”

Steinhauer, K., K. Alter and A. D. Friederici (1999). “Brain potentials indicate immediate use of prosodic cues in natural speech processing.” Nature neuroscience 2(2): 191.

Sun, Y., F. Gomez and J. Schmidhuber (2011). Planning to Be Surprised: Optimal Bayesian Exploration in Dynamic Environments. Artificial General Intelligence: 4th International Conference, AGI 2011, Mountain View, CA, USA, August 3-6, 2011. Proceedings. J. Schmidhuber, K. R. Thórisson and M. Looks. Berlin, Heidelberg, Springer Berlin Heidelberg:41–51.

Sur, M., P. E. Garraghty and A. W. Roe (1988). “Experimentally induced visual projections into auditory thalamus and cortex.” Science 242(4884): 1437–1441.

Taylor, J. S., K. Rastle and M. H. Davis (2013). “Can cognitive models explain brain activation during word and pseudoword reading? A meta-analysis of 36 neuroimaging studies.” Psychol Bull 139(4): 766–791.

Tervaniemi, M., T. Ilvonen, K. Karma, K. Alho and R. Näätänen (1997). “The musical brain: brain waves reveal the neurophysiological basis of musicality in human subjects.” Neuroscience letters 226(1): 1–4.

Tervaniemi, M., I. Winkler and R. Näätänen (1997). “Pre-attentive categorization of sounds by timbre as revealed by event-related potentials.” NeuroReport 8(11): 2571–2574.

Thiel, A., B. Habedank, L. Winhuisen, K. Herholz, J. Kessler, W. F. Haupt and W. D. Heiss (2005). “Essential language function of the right hemisphere in brain tumor patients.” Ann Neurol 57(1): 128–131.

Thiessen, E. and L. Erickson (2013). “Discovering Words in Fluent Speech: The Contribution of Two Kinds of Statistical Information.” Frontiers in Psychology 3(590).

Toiviainen, P., M. Tervaniemi, J. Louhivuori, M. Saher, M. Huotilainen and R. Näätänen (1998). “Timbre similarity: Convergence of neural, behavioral, and computational approaches.” Music Perception: An Interdisciplinary Journal 16(2): 223–241.

Tourville, J. A. and F. H. Guenther (2011). “The DIVA model: A neural theory of speech acquisition and production.” Lang Cogn Process 26(7): 952–981.

Ueno, T., S. Saito, T. T. Rogers and M. A. Lambon Ralph (2011). “Lichtheim 2: synthesizing aphasia and the neural basis of language in a neurocomputational model of the dual dorsal-ventral language pathways.” Neuron 72(2): 385396.

Ulanovsky, N. and C. F. Moss (2008). “What the bat’s voice tells the bat’s brain.” Proceedings of the National Academy of Sciences of the United States of America 105(25): 8491–8498.

Ungerleider, L. G. and J. V. Haxby (1994). “‘What’ and ‘where’ in the human brain.” Current Opinion in Neurobiology 4(2): 157–165.

Van Dommelen, W. A. (1987). “The Contribution of Speech Rhythm and Pitch to Speaker Recognition.” Language and Speech 30(4): 325–338.

Van Dommelen, W. A. (1990). “Acoustic parameters in human speaker recognition.” Language and Speech 33(3): 259–272.

Van Petten, C., S. Coulson, S. Rubin, E. Plante and M. Parks (1999). “Time course of word identification and semantic integration in spoken language.” Journal of Experimental Psychology: Learning, Memory, and Cognition 25(2): 394.

Van Petten, C. and M. Kutas (1990). “Interactions between sentence context and word frequencyinevent-related brainpotentials.” Memory & cognition 18(4): 380–393.

Vanthornhout, J., L. Decruy, J. Wouters, J. Simon and T. Francart (2018). “Speech intelligibility predicted from neural entrainment of the speech envelope.” bioRxiv(637424): 246660.

Veale, R., Z. M. Hafed and M. Yoshida (2017). “How is visual salience computed in the brain? Insights from behaviour, neurobiology and modelling.” 372(1714).

Vinckier, F., S. Dehaene, A. Jobert, J. P. Dubus, M. Sigman and L. Cohen (2007). “Hierarchical coding of letter strings in the ventral stream: Dissecting the inner organization of the visual word-form system.” Neuron 55(1): 143–156.

Wacongne, C., J. P. Changeux and S. Dehaene (2012). “A neuronal model of predictive coding accounting for the mismatch negativity.” J Neurosci 32(11): 3665–3678.

Walden, B. E., A. A. Montgomery, G. J. Gibeily, R. A. Prosek and D. M. Schwartz (1978). “Correlates of psychological dimensions in talker similarity.” Journal of speech, language, and hearing research 21: 265–-275.

Warburton, E., C. J. Price, K. Swinburn and R. J. S. Wise (1999). “Mechanisms of recovery from aphasia: evidence from positron emission tomography studies.” Journal of Neurology, Neurosurgery & Psychiatry 66(2): 155–161.

Winkler, I., S. L. Denham and I. Nelken (2009). “Modeling the auditory scene: predictive regularity representations and perceptual objects.” Trends in Cognitive Sciences 13(12): 532–-540.

Winn, J. and C. M. Bishop (2005). “Variational message passing.” Journal of Machine Learning Research 6: 661–694.

Ylinen, S., M. Huuskonen, K. Mikkola, E. Saure, T. Sinkkonen and P. Paavilainen (2016). “Predictive coding of phonological rules in auditory cortex: A mismatch negativity study.” Brain Lang 162: 72–80.

Zeki, S. and S. Shipp (1988). “The functional logic of cortical connections.” Nature 335: 311–317.

Zhang, C., J. Butepage, H. Kjellstrom and S. Mandt (2018). “Advances in Variational Inference.” IEEE Trans Pattern Anal Mach Intell.

